# HM-DyadCap – Capture and Mapping of 5-Hydroxymethylcytosine/5-Methylcytosine CpG Dyads in Mammalian DNA

**DOI:** 10.1101/2025.10.29.685270

**Authors:** Lena Engelhard, Damian Schiller, Marlon S. Zambrano-Mila, Kotryna Keliuotyte, Benjamin Buchmuller, Shashank Tiwari, Jochen Imig, Angela Simeone, Christian Schröter, Sidney Becker, Daniel Summerer

**Affiliations:** Faculty of Chemistry and Chemical Biology, TU Dortmund University, Otto-Hahn-Str. 4a, 44227 Dortmund (Germany); Max-Planck-Institute for Molecular Physiology, Otto-Hahn-Str. 11, 44227 Dortmund (Germany); Genomic Services, Qiagen, Manchester, UK

**Author notes:** these authors contributed equally to this work (Lena Engelhard, Damian Schiller).

## Abstract

5-Methylcytosine (mC) and 5-hydroxymethylcytosine (hmC) are the main epigenetic modifications of mammalian DNA, and play crucial roles in cell differentiation, development, and tumorigenesis. Both modifications co-exist with unmodified cytosine in palindromic CpG dyads in different symmetric and asymmetric combinations across the two DNA strands, each having unique regulatory potential. To facilitate investigating the individual functions of such dyad modifications, we report HM-DyadCap. This method employs an evolved methyl-CpG-binding domain (MECP2 HM) for the direct capture and sequencing of DNA fragments containing the CpG dyad hmC/mC. Binding studies reveal a high discrimination of MECP2 HM against off-target dinucleotides. We conduct comparative mapping experiments for mESC genomes with HM-DyadCap, standard MethylCap employing wild type MECP2, as well as MeDIP and hMeDIP protocols. We find that MECP2 HM is blocked by hmC glucosylation, and conduct control enrichments with glucosylated genomes that indicate highly selective enrichment of hmC/mC dyads by MECP2 HM. Metagene profiles correlate hmC/mC marks with actively transcribed genes, and reveal global enrichment in gene bodies as well as depletion at transcription start sites. We anticipate that HM-DyadCap will enable effective enrichment and mapping of hmC/mC marks with broad applicability for unravelling the function of this dyad in chromatin biology and cancer.

## INTRODUCTION

The epigenetic DNA modification 5-methylcytosine (mC, **Fig. 1a**) is a central regulator of mammalian gene expression with crucial roles in development, differentiation, and cancer formation^1^. mC is written and maintained over cell cycles by DNA methyltransferases (DNMTs) mainly within palindromic CpG dyads, of which 60-80 % are methylated in somatic cells^2^. mC can be read and converted into transcriptionally repressive states by methyl-CpG-binding domain (MBD) proteins^3^, but can also recruit or repel transcription factors and other chromatin proteins^4^. mC can further be oxidized by ten-eleven translocation (TET) dioxygenases, which generate CpG dyads containing 5-hydroxymethylcytosine (hmC, **Fig. 1a**), 5-formylcytosine (fC) and 5-carboxylcytosine (caC). Whereas fC and caC are substrates of the thymine-DNA glycosylase (TDG)-initiated base excision repair (BER) pathway leading to an active demethylation^5,6^, hmC exhibits high stability, and occurs at high levels in embryonic stem cells (ESC) and neurons^5–7^. hmC thereby differs from mC in its physicochemical properties, genomic distributions, dynamic changes during development, and protein interactions, and thus has potential to uniquely regulate chromatin-associated processes^5,8,9^.

**Figure 1.**
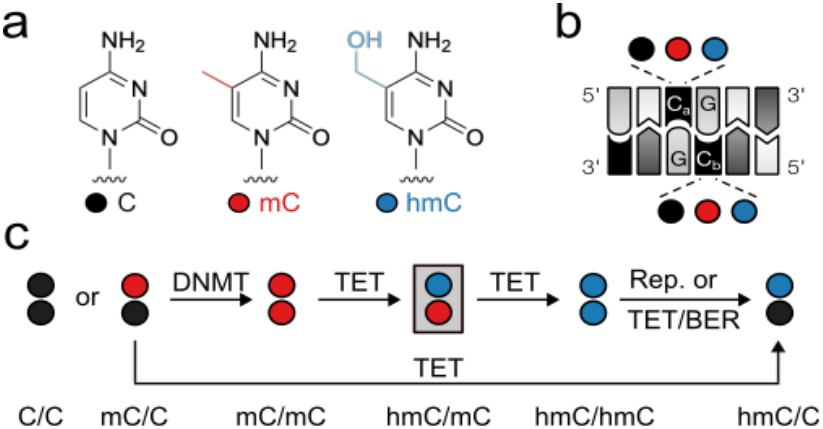
Mammalian cytosine modifications in the double-stranded CpG dyad. a) Structures of C, mC, hmC (fC and caC not shown). b) Both cytosines (C_a_, C_b_) in the CpG dyad can exist as C, mC or hmC. c) Possible pathways for the creation of different hmC dyad symmetries (hmC/mC dyad targeted in this study in grey box) by DNMTs, TETs, replication (Rep.) and BER.

For example, whereas hmC is enriched in promoters and gene bodies in mouse ESC (mESC) and neurons^10–12^, it shows generally low levels and different genomic distributions in cancer cells, making it an important cancer biomarker^13,14^.

In contrast to mC that can be kept in a strand-symmetric state in CpGs by maintenance DNMTs, the writing of hmC and other oxi-mCs by TETs occurs non-processively^15^, leading to different strand-symmetric and -asymmetric combinations of C, mC and hmC in the CpG dyad (**Fig. 1b**). The genomic landscape of these marks can further be shaped by BER, the inhibition of maintenance methylation^5^, and TET-catalyzed oxidation of hemimethylated mC/C dyads^16^ (**Fig. 1c**). Each of the aforementioned dyads presents a physicochemically unique signal in the DNA major groove^17^, an important interaction surface for DNA binding proteins^8^. The question of how hmC has unique regulatory functions can thus only be answered conclusively by considering the hmC-modification symmetry of CpG dyads.

Sequencing and mapping studies of hmC have greatly contributed to a better understanding of its functions, but established mapping methods do not simultaneously resolve C, mC and hmC in the same DNA duplexes^5,18^. Very recently, new strategies for the simultaneous sequencing of C, mC and hmC have been reported, offering potential for refined maps with resolution for individual hmC dyad symmetries. These employ protocols based on restriction enzymes (DARESOME^19^ and Dyad-seq^20^), on multiple consecutive nucleobase conversions to achieve nucleotide resolution (e.g., EnIGMA^21^, SCoTCH-Seq^22,23^, and SIMPLE-Seq^24^), or on direct nanopore sequencing^25^. In two cases, strategies have been adapted/applied for mapping individual hmC dyads. This revealed that in mESC and mouse cerebellum genomes, the large majority of hmC resides in asymmetric hmC/mC dyads, whereas hmC/C and symmetric hmC/hmC dyads are comparably rare^20,25^. A third, very recent mESC genome mapping study using SCoTCH-seq reported frequencies in the order hmC/C>hmC/mC>>hmC/hmC^23^.

A deeper understanding of the individual hmC dyad’s roles in chromatin regulation during development and disease requires broader mapping studies across diverse tissues, which however are complicated by the low hmC levels found in most tissues (e.g., cancer tissues)^6,7^. Methods for the effective enrichment of hmC-modified DNA could resolve this bottleneck by direct sequencing/mapping, or in combination with aforementioned conversion-based sequencing. However, current hmC-enrichment strategies rely on anti-hmC antibodies^26–28^ or T4-β-glucosyltransferase (T4 BGT)-catalyzed azidoglucose transfer to hmC and covalent capture by click chemistry^29,30^, both of which have not been reported to be dyad-specific^31^ (e.g., T4 BGT effectively glucosylates hmC in different dyad contexts, **Fig. S1**).

To address this bottleneck, we developed HM-DyadCap, a method that employs the evolved MECP2 variant MECP2 HM for selective enrichment and sequencing of hmC/mC CpG dyads that are abundant marks mESC and brain genomes. In vitro binding studies show that MECP2 HM has a high specificity for its target dyad. We conduct comparative mapping studies with HM-DyadCap, MethylCap based on MECP2 wild type (MECP2 wt), as well as antibody-based methyl- and hydroxymethylcytosine-DNA-immunoprecipitation protocols (MeDIP and hMeDIP), and observe method-dependent preferences in the enrichment of specific genomic features. We discover a sensitivity of MECP2 HM to hmC glucosylation and introduce additional mapping controls with glucosylated genomes that indicate a high hmC/mC selectivity of HM-DyadCap. Metagene profiles reveal enrichment of hmC/mC in gene bodies and depletion at transcription start sites, and correlate it with active transcription. HM-DyadCap offers a simple and effective approach for enriching and mapping hmC/mC dyads, and we anticipate broad applicability for cancer biomarker discovery and unravelling the function of hmC/mC in chromatin regulation.^64^

## MATERIAL AND METHODS

### mESC culturing and gDNA isolation

E14tg2a mouse embryonic stem cells^32^ were grown in dishes pre-coated with 0.1 % gelatin (w/vol, Sigma) in GMEM supplemented with 10% fetal bovine serum, sodium pyruvate, 50 μM β-mercaptoethanol, glutamax, non-essential amino acids (all from Gibco/ThermoFisher) and 10 ng/ml murine leukemia inhibitory factor (LIF, Protein Expression Facility, MPI Dortmund). Cells were passaged every 2-3 days to maintain cultures between 10% and 90% confluency. For analysis, near-confluent cultures were released from culture vessels with trypsin, spun down, and snap-frozen in liquid nitrogen before further processing. gDNA was isolated using the Monarch Genomic DNA Purification Kit (NEB, T3010) according to the manufacturer’s instructions. DNA concentration and purity were assessed using Nanodrop.

### gDNA fragmentation

gDNA at a concentration of 10 ng/µl in 1xTE was sheared to an average fragment size of 200 bp using a Bioruptor Pico sonication device (Diagenode). Fragmentation was performed for 19 cycles of 30 s on, 30 s off at 4 °C. Sheared DNA was purified by ethanol precipitation overnight at −80 °C. dsDNA concentration was measured with a Quantus Fluorometer (Promega). Fragment size distribution was confirmed using a TapeStation (Agilent Technologies) using D1000 ScreenTapes. Sheared DNA was either subjected to glucosylation or directly used for Illumina library preparation.

### Glucosylation of 5hmC

For each glucosylation reaction, 1 µg of sheared gDNA was used. Reactions were carried out in 1x Cutsmart buffer (NEB) supplemented with 40 µM UDP-Glucose and 20 U T4-BGT (NEB, M0357) in a total volume of 40 µl for 16 h at 37 °C. Subsequently, DNA was purified using the Monarch PCR & DNA Purification kit following the manufacturers protocol for fragments < 2 kbp. DNA concentration was measured with a Quantus Fluorometer (Promega). For glucosylated hmC binding studies, 500 ng of 24-mer duplex DNA (hmC/mC: o2909 + o3115; hmC/hmC: o2909 + o3115, **Table S1**) was incubated with 10 U T4-BGT (NEB) under conditions described above. Subsequently, DNA was purified using the Oligo Clean and Concentrator kit (Zymo Research) following the manufacturers protocol.

### End repair and adapter ligation

1 µg fragmented DNA, either glucosylated or non-glucosylated, was subjected to end repair and adapter ligation using the NEBNext Ultra II DNA Library Preparation Kit (NEB, E7103; version 6.1_5/20). DNA purification and size selection was carried out using NEBNext Sample Purification Beads (NEB, E7103) according to the protocol with bead volumes corresponding to a target fragment size of 200 bp. Elution was carried out according to the manufacturers protocol, and DNA concentration was determined with the Quantus Fluorometer (Promega). Fragment size distribution was verified using a TapeStation (Agilent Technologies) with D1000 ScreenTapes.

### Spike-in probe preparation

Preparation of spike-in probes was carried out as described previously^33^. In brief, DNA duplexes carrying modified CpG dyads (“carriers”) were ligated to adapter sequences containing general and unique primer binding sites for qPCR quantitation. Carriers were prepared by annealing modified oligonucleotides (C/C: o4728 + o4729, mC/mC: o4677 + o4727, hmC/mC: o4675 + o4727) in 30 mM HEPES pH 7.5 and 100 mM KOAc. Annealing was carried out by heating to 95 °C for 5 min followed by gradual cooling in a Dewar flask containing boiling water. Adapters were annealed at a final concentration of 2.5 µM in the same way (o4371 + o4372, o4373 + o4374, and o4392 + o4393) and subsequently extended with 25 mU/µl Klenow fragment (NEB) in the presence of 0.1 mM dNTPs for 20 min at 37 °C. To introduce 5’ phosphorylation, primer o4123 was annealed to the extended adapters and further extended with Klenow fragment for 30 min at 37 °C. 50 µl of crude adapters were ligated with15 pmol of carriers at 16 °C overnight using 200 U T4 DNA ligase (NEB). Ligated spike-in probes were purified using the Macherey-Nagel PCR purification kit, with the NTI buffer diluted to 16 % (v/v) with ddH_2_O. Probe concentration was determined by qPCR against reference standards (o4628 for o4374, o4627 for o4372 and o4629 for o4393). qPCR primers o4368 and o4124 were premixed as a 2 µM stock. Each reaction contained a 3 µl primer mix, 5 µl 2x primaQuant SYBRGreen Master Mix with ROX (Steinbrenner) and 2 µl of spike-in probe. Cycling conditions were 95 °C for 2 min, followed by 50 cycles of 95 °C for 10 s and 60 °C for 30 s.

### MECP2 wt and MECP2 HM expression and purification

GST-tagged MBP-MECP2 wt and MBP-MECP2 HM were recombinantly expressed in *E. coli* BL21-Gold (DE3). A 300 ml LB culture supplemented with 1 mM MgCl_2_, 1 mM ZnSO_4_ and a suitable antibiotic was inoculated from a fresh overnight culture and grown at 37 °C with shaking at 220 rpm to an OD_600_ of 0.5–0. 6. Cultures were chilled on ice and protein expression was induced by the addition of 1 mM IPTG. The expression proceeded overnight at 30 °C with shaking at 150 rpm. The cultures were harvested at 8,000 x g for 20 min at 4 °C, washed twice by resuspension in 50 ml cold 20 mM Tris-HCl (pH 8.0), and the resulting pellet was frozen until further processing. For lysis, cell pellets were resuspended in 20 ml binding buffer (20 mM Tris-HCl, 250 mM NaCl, 10% glycerol, 10 mM dithiothreitol (DTT), 5 mM imidazole, 0.1% Triton X-100, pH 8.0) supplemented with 1 mM PMSF. The suspension was treated with 0.1 mg/ml lysozyme (Merck) and 1 U/ml DNase I (NEB) and incubated overnight at 4 °C. Cells were disrupted by two rounds of sonication on ice (3 min per run of alternating a 4 s ultrasonic wave pulse at 20 % amplitude and 30 s rest (Branson Digital Sonifier 450 Cell Disruptor). Cellular debris was removed by centrifugation at 14,000 x g for 20 min at 4 °C and the cleared supernatant was retained and purified over a Ni-NTA column (GE Healthcare) on an Äkta Purifier 10 FPLC system (GE Healthcare) using a gradient of imidazole (10 mM to 500 mM) in binding buffer. The fractions that contained the pure protein were combined and dialyzed three times against dialysis buffer (20 mM HEPES, 100 mM NaCl, 10 % glycerol, adjusted to pH = 7.3, and 0.1 % Triton X-100) using Slide-A-Lyzer dialysis cassettes (3.5 kDa MWCO, ThermoFisher Scientific). The protein concentration was determined in triplicate using the Pierce BCA Protein Assay (ThermoFisher Scientific). Proteins were snap frozen in liquid nitrogen and stored at –80 °C at a concentration of 15 µM.

### Electrophoretic mobility shift assays

Complementary oligonucleotide pairs (**Table S1**) were annealed by mixing 1.5 μM of the FAM-labeled strand and 2.5 μM of the unlabeled strand in EMSA hybridization buffer (20 mM HEPES, 30 mM KCl, 1 mM EDTA, 1 mM (NH_4_)_2_SO_4_, pH 7.3), incubated at 95 °C for 5 min, and gradually cooled down to RT in a Dewar filled with boiling water. The non-specific dA:dT competitor duplex was prepared by annealing 24-mer poly(A) with poly(T) oligos (o2968 + o2969, **Table S1**) at an equimolar ratio of 40 μM. Prior to the binding assay, 15 μM of purified recombinant proteins were incubated with 0.25 μM TEV protease at 4 °C overnight to remove the MBP tag. For glucosylated hmC binding studies, varying concentrations of protein were incubated with 2 nM labeled duplex DNA in EMSA buffer (20 mM HEPES, 30 mM KCl, 1 mM EDTA, 1 mM (NH_4_)_2_SO_4_, 0.2 % Tween-20, pH 7.3) in presence of 3.4 µM poly(A):poly(T) competitor and 1 mM DTT in a total volume of 15 µL. Reactions were incubated for 20 min at 21 °C. For assays using DNA probes with CpG in random sequence contexts, 100 nM of protein was incubated with 2 nM labeled probe under the same binding conditions. For antibody binding assays, 750 pM of labeled DNA probes were incubated with either polyclonal anti-5hmC antibody (Active Motif, 39769; 1:100 and 1:200 dilutions, **Fig. 4b-c**), monoclonal anti-5hmC antibodies (EpiGentek (EG), A-1018; Cell Signaling (CS), 51660; Diagenode (DG), C15200200; each at 1:10 dilutions, **Fig. 4a**), or 100 nM MECP2 HM protein (HM in **Fig. 4a**) in binding buffer (10 mM Tris, 100 mM KCl, 1 mM EDTA, 5 % glycerol, 0.1 mg/ml BSA, pH 7.5) containing 3.4 µM poly(A):poly(T) competitor in a total volume of 15 µL and incubated at 4 °C overnight. After incubation, samples were mixed with 3 µL of 6x EMSA loading buffer (1.5x TBE, 40 % glycerol) and resolved on pre-run 15 % non-denaturing polyacrylamide gels. Electrophoresis was performed at 240 V for 45 min (MECP2 binding assays) or 55 min (antibody binding assays) at 4 °C using Mini-PROTEAN vertical electrophoresis units (Bio-Rad). Gel fluorescence was recorded using the 510 LP filter of a Typhoon FLA9500 laser scanner (GE Healthcare) at 473 nm at 800 - 1000 V amplification. The fraction of bound dsDNA was determined using ImageQuant TL v8.1 1D Gel Analysis (DE Healthcare), applying background subtraction and manual peak detection with approximately equal peak areas across all lanes.

### Mass spectrometry

20 µl of 0.1 µM DNA sample was analyzed per injection in the Vanquish HPLC system connected to an Orbitrap Exploris 120 ESI-MS (Thermo Fisher Scientific). The sample components are separated with a flow rate of 0.2 ml/min in a solvent system of 50 mM HFIP (1,1,1,3,3,3-Hexafluoro-2-propanol, 15 mM triethylamine, pH = 9.0 (solvent A), and methanol (solvent B) at 80 °C on DNAPac RP 4 µm column (2.1 × 100 mm, Thermo Scientific). A multistep gradient (3 % of solvent B in 0–2 min, followed by an increase from 3 to 10 % until 5 min and 10 to 20 % until 20 min) was used for separation. MS analysis was performed in the negative mode with a spray voltage of 2500 V, flow of sheath gas, aux gas, and sweep gas maintained at 35, 7, and 0, respectively, and ion transfer tube temperature and vaporizer temperature set at 300 °C and 275 °C, respectively. For MS1 analysis, the Orbitrap resolution of 120,000 was used, the scan range was set from 700 to 3000 m/z, RF lens at 70 %, with standard normalized AGC target and data type set to profile mode. All MS data analysis was performed with Biopharma Finder v.05.1, automatic parameter values were set for component detection. For oligonucleotide identification, output mass range was set from 2 kDa to 12 kDa, a charge range of 3-20 were considered with a minimum detected charge of 3. Of the identified fragments, only those with a mass error less than 10 ppm were considered for further analysis.

### Genomic enrichment of modified CpG dyads

For enrichment, an equimolar mix of spike-in probes (6.04 fmol each) was added to 250 ng of sheared gDNA in Buffer B (from MethylCap Kit, Diagenode) in a total volume of 35.45 µl. From this mixture, 5.7 µl were set aside as input and the remaining 29.75 µl were used for the enrichment reaction. GST-tagged MBP-MECP2 wt and MBP-MECP2 HM were TEV-digested for 30 min at RT to remove MBP. TEV-digested proteins were then added to the DNA mixture at a final concentration of 1.7 µM in a total reaction volume of 34 µl. Enrichment was performed according to the manufacturer’s instructions (MethylCap Kit Diagenode, version 6, C02020010) following the high-salt elution protocol. MECP2 wt and MECP2 HM enrichments were carried out in separate reactions, each performed in three technical replicates per biological experiment. Eluted DNA was purified using the Macherey Nagel PCR purification kit. Spike-in probe recovery was determined by qPCR against a standard dilution series of spike-in probes. To distinguish individual spike-ins, three different primer pairs were used (C/C: o4374+o4368, mC/mC: o4372+o4368, hmC/mC: o4393+o4368). qPCR was performed on the elution fractions. Reactions were carried out by initial denaturation at 95 °C for 2 min, followed by 50 cycles of 95 °C for 10 s and 60 °C for 30 s. The remaining elution samples were further processed and prepared for NGS.

### Preparation of Illumina libraries and NGS sequencing

Enriched DNA fragments were PCR amplified using the NEBNext Ultra II DNA Library Prep Kit for Illumina (NEB, version 6.1_5/20) using NEBNext Multiplex Oligos (NEB, E73359). As amplification does not occur directly after adapter ligation but after the enrichment, DNA input concentrations were treated as threefold higher for determining the appropriate number of PCR cycles. PCR clean-up was performed according to the “Cleanup of PCR Reaction” protocol (version 6.1_5/20) using NEBNext Sample Purification Beads. Fragment length distribution of the final libraries was assessed with a Tapestation (Agilent Technologies) with a D1000 ScreenTape according to the Agilent protocol. dsDNA concentration was measured with the Quantus Fluorometer (Promega). Libraries were pooled in equimolar ratios and sequenced on a NovaSeq X-25B instrument (Illumina) using paired-end reads, with a target depth of 50 million reads per sample.

### MeDIP and hMeDIP

For each immunoprecipitation, 1 µg of sheared gDNA was used as input. DIPs were performed according to the MagMeDIP-seq Package V2 protocol (Diagenode, C02010041), using either the provided anti-5mC antibody for MeDIP or 0.6 µg per IP of the mouse monoclonal anti-hmC antibody (Diagenode, C15200200) for hMeDIP. Additionally, a control IP with 0.6 µg of mouse IgG (Diagenode, C15400001) was included. Illumina libraries were prepared according to the Diagenode MeDIP-seq library preparation protocol. Final library concentrations were quantified using the Quantus Fluorometer (Promega) and fragment size distributions were assessed with a TapeStation (Agilent Technologies) using a D1000 ScreenTape. Libraries were sequenced on an Illumina NovaSeq X-25B instrument using paired-end 2x 150 bp reads with a target depth of 50 million reads per sample.

### Data Analysis

Paired-end sequencing libraries were processed using a standardized and automated workflow implemented with Snakemake^34^. Initial quality assessment of raw reads was carried out using FastQC^35^, followed by adapter trimming and removal of low-quality bases using Trim Galore^36^. The cleaned reads were then aligned to the *Mus musculus* reference genome mm10 using Bowtie2^37^. Retaining only properly paired reads and removing PCR duplicates was done using Samtools^38^. To identify enriched regions, peak calling was performed on deduplicated BAM files using MACS2^39^. Technical reproducibility was assessed by conducting peak calling independently for each of the three technical replicates along with their respective input controls within each biological replicate. Reproducibility was evaluated across three independent replicates by generating a consensus peak set, retaining peaks present in at least two out of the three replicates using bedtools^40^. In the case of DIP-seq, enriched regions were obtained as previously described using MACS2, with either IgG or input from mESCs serving as controls. True positive regions were defined as those enriched regions identified for both IgG and input controls, as described by Lentini et al., 2018. For signal visualization, normalized BigWig files were generated using bamCoverage (deepTools v3.5.0^41^), and final alignments were visually inspected in the Integrative Genomics Viewer (IGV^42^). To assess the statistical significance of overlap between consensus peaks and annotated genomic features, the Genomic Association Tester (GAT^43^) was employed. Metagene profiles were generated to assess enrichment relative to gene expression. Log_2_ fold changes over input were calculated across gene bodies using bamCompare (deepTools). Genes were ranked by expression, defined as the geometric mean of FPKM from RNA-seq (E14 ENCODE replicates 1 and 2), and grouped into highly expressed (top 10%), lowly expressed (FPKM > 0.1 and below the top 10% threshold), and silenced (FPKM < 0.1). Profiles were computed with computeMatrix (deepTools) and visualized using plotHeatmap (deepTools). Additional downstream analyses were performed using bedtools, deepTools, and custom Bash and R scripts.

## RESULTS AND DISCUSSION

### Selectivity of MECP2 wt and MECP2 HM in respect to CpG dyad-specific genomic enrichment

Enrichment of genomic DNA fragments bearing specific epigenetic modifications greatly increase the efficiency of mapping studies by overcoming the need for whole genome sequencing. For hmC, antibodies and T4 BGT are both broadly used enrichment tools regardless of the specific CpG dyad symmetry. While MBD proteins are also popular probes for genomic enrichment, they specifically recognize symmetrically methylated (mC/mC) CpG dyads in native dsDNA, and are therefore limited to mC enrichment assays^44^. Indeed, the functional members of mammalian core family MBD proteins (MBD1, 2, 4 and MECP2) tend to be repelled by hmC^45–49^. To expand the scope of MBD-based technologies, we recently re-evolved the MBD domain of human MECP2 wt to switch its selectivity from the canonical mC/mC CpG to TET-generated CpG dyads^33,50^. In this course, we identified the mutant MBD domain MECP2 HM that selectively recognizes the dyad hmC/mC (mutations K109T, V122A, S134N^51^). In electromobility shift assays (EMSA), this mutant showed a similar on-versus main off-target selectivity (hmC/mC over mC/mC) as MECP2 wt (mC/mC over hmC/mC). Moreover, it exhibited a higher discrimination as MECP2 wt against all other CpG dyads containing C, mC or hmC, particularly against the frequent dyad modifications mC/C and hmC/C^23^ (**Fig. 2a**, data from ref^33^). Importantly, MECP2 HM is based on a minimal MBD domain of MECP2 (aa 90-181 of the 486 aa full-length protein) that does not contain AT-hook sequences, which have been shown to cause a weak preference for AT-rich sequence contexts around the target CpG^52,53^. Context-dependence has also been described for antibody-based DIP-Seq protocols that suffer from significant nonspecific enrichment of short repeat sequences via IgG^54–56^, a property that has not been observed in previous enrichments using the MBD of MECP2 wt^56^.

**Figure 2.**
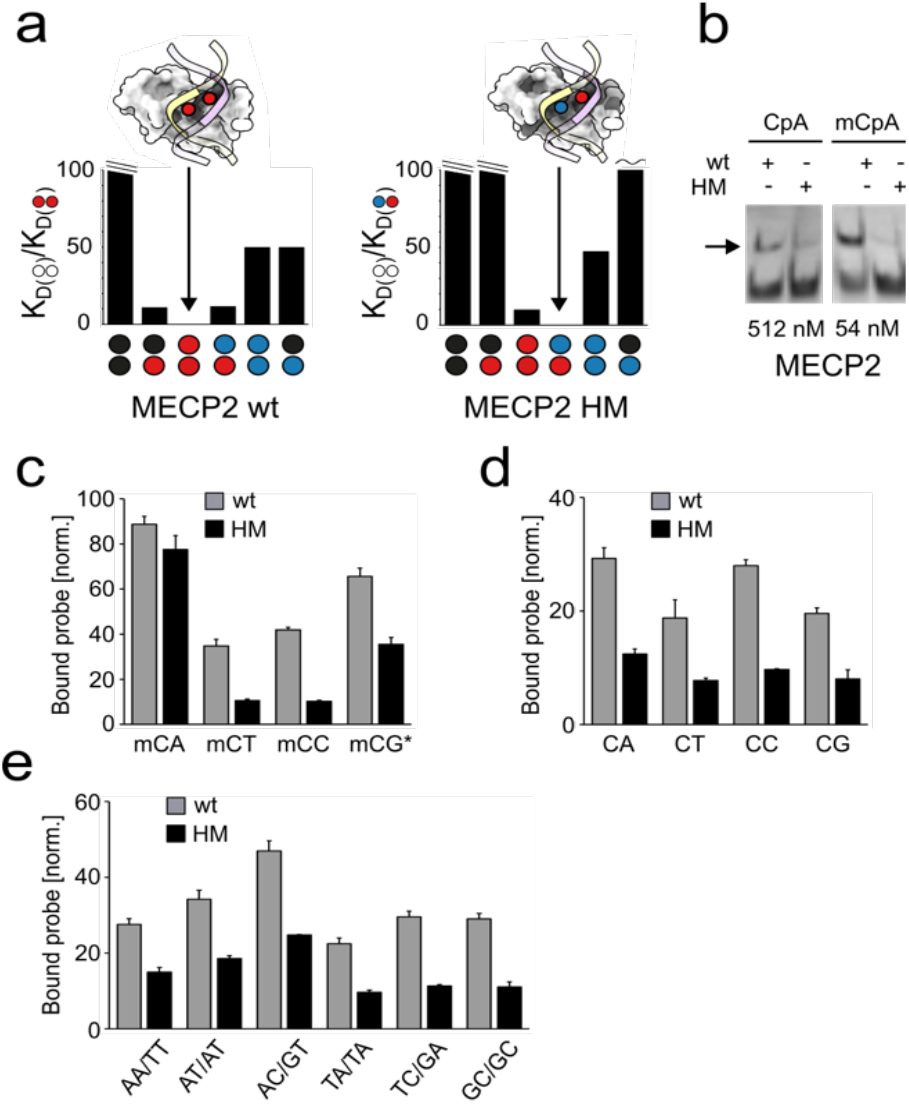
Characterization of off-target dinucleotide selectivity of MECP2 wt and HM. a) Selectivity of MECP2 wt and HM for indicated CpG modification symmetries. Shown are off-to on-target K_D_ ratios from EMSA using dsDNA probes with a single CpG in an oligo dA/dT context (data from ref^33^, color code as in **Fig. 1a**). b) EMSA with MECP2 wt and HM at indicated concentrations and 2 nM dsDNA probes with single CpA or mCpA in an oligo dA/dT context (arrow: bound probe). c-e) Quantification of EMSA with MECP2 wt or HM and dsDNA probes with indicated methylated or unmethylated dinucleotides in a random sequence context (*hemi-methylated CpG). Shown are the % fractions of protein-bound probe normalized to the respective on-target of MECP2 wt (mC/mC) and HM (hmC/mC). Error bars show standard deviation from n=2.

To further characterize the selectivity of MECP2 HM in respect to relevant genomic off-targets, we initially tested its interaction with mCpA dinucleotides by EMSA. mCpA occurs at significant levels in neuronal and embryonic stem cells^57,58^, and it is bound by MECP2 wt (aa 1-205)^59^. Interestingly, we observed significantly lower binding of MECP2 HM to both methylated and non-methylated CpA as compared to MECP2 wt when present in an oligo dA/dT sequence context (**Fig. 2b**). Encouraged by this finding, we conducted broader EMSA studies covering other possible off-target dinucleotides. MECP2 wt has been shown to bind mCpA with a strong preference for a 3’-A nucleotide (mCAA; a property not observed for CpG dyads^60^). To rule out such possible context dependencies, we employed random dsDNA probes in EMSA (5’-…NNNXYNNN…-3’). Finally, in order to visualize binding to even very low affinity off-targets, we applied a high, 50-fold excess of MECP2 over DNA. As expected, we observed mCpA off-target binding for both proteins under these forcing conditions (**Fig. 2c** and **S2**). Hemi-methylated mCpG dyads were bound with slightly lower and mCpT as well as mCpC with much lower relative affinity, in agreement with previous studies^60^. Interestingly, MECP2 HM showed higher selectivity over off-targets in all cases (**Fig. 2c** and **S2**). We next tested nonmethylated CpN dinucleotides and observed a generally much lower binding by both proteins, with a different selectivity profile as observed for mCpN (**Fig. 2d** and **S2**). Strikingly, CpN off-targets were again bound weaker by the HM mutant as compared to the wt protein. For a comprehensive analysis of the so far uncharacterized dinucleotide selectivity of our evolved MECP2 HM, we finally tested probes containing all remaining, nonmethylated NpN dinucleotides.

We observed a similar trend with particularly low relative off-target binding for MECP2 HM, corroborating the CpG-selectivity of this protein (**Fig. 2e** and **S2**). In the light that MECP2 wt is a widely used enrichment probe for mC/mC CpGs, the observed high selectivity of MECP2 HM underlines its suitability for enrichments of genomic hmC/mC CpG dyads with little potential for off-target enrichment.

We next established a simple protocol for enriching hmC/mC-modified DNA fragments from mammalian genomes (**Fig. 3a**). Briefly, gDNA is sheared via sonication and subjected to end-repair and adaptor ligation for high throughput sequencing (**Fig. 3a** and **S3-6**). After purification, three spike-in control DNAs containing a sequence with four CpG dyads (either unmodified, mC/mC- or hmC/mC-modified) are added in equimolar amounts.

**Figure 3.**
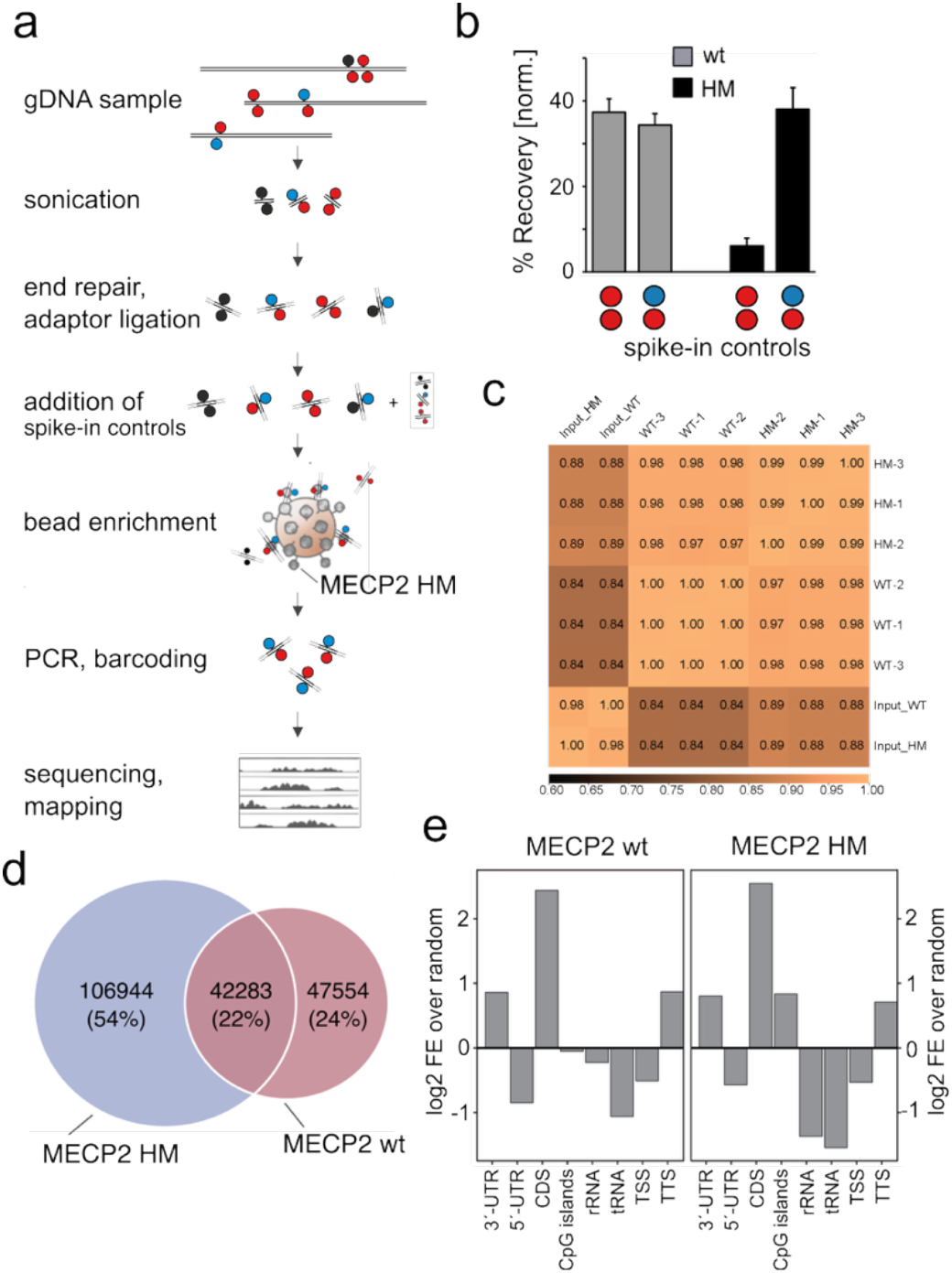
Enrichment and sequencing of modified CpG from mESC genomes with MECP2 wt and HM. a) Workflow of HM-DyadCap. b) Recovery of spike-in controls from genomic enrichments. c) Spearman correlation of triplicate enrichment conditions. d) Venn diagrams of peaks obtained from enrichments with MECP2 wt and HM. e) Enriched or depleted genomic features from enrichments with MECP2 wt and HM.

Each of these controls contains unique primer binding sites for selective quantification by qPCR, but lack adaptors to not interfere with downstream sequencing. For each enrichment, 250 ng gDNA are incubated with MECP2 HM-GST fusion protein immobilized on glutathione magnetic beads, and the DNA is eluted after washing. The samples are subjected to adaptor PCR with sample barcoding and then purified and sequenced (**Fig. 3a**). Fragment size distribution is controlled after adapter ligation and the final PCR, and enrichment success is assessed by qPCR-measurement of mC/mC and hmC/mC spike-in recoveries normalized to the unmodified spike-in control directly after bead enrichment (**Fig. 3b**). To generate enrichment-based maps of hmC/mC CpG dyads in a mammalian genome, we conducted enrichments with MECP2 HM and MECP2 wt using mESC genomic DNA (E14tg2a cell line), a system where epigenetic regulation is particularly essential.

Using spike-in oligonucleotides, we assessed the dyad selectivity of both MECP2 probes under enrichment conditions. Recovery of spike-ins showed low specificity for MECP2 wt (i.e., enrichment of both mC/mC and hmC/mC), whereas MECP2 HM selectively enriched its hmC/mC target (**Fig. 3b**)^33^. Sequencing of these libraries produced between ~63 million and 184 million reads (**Table S2**), with over 95% aligning to the mm10 mouse reference genome. To assess the enrichment efficiency, we compared the percentage of reads within MACS2-called peaks between input and captured samples across MECP2 conditions (**Fig. S7**). In both cases, captured samples showed higher enrichment than their respective inputs, with the stronger effect observed for MECP2-HM. These results confirm the robust enrichment achieved with our protocol.

To evaluate reproducibility and overall similarity across datasets, we calculated Spearman correlations across 100 kb genomic bins and performed hierarchical clustering (**Fig. 3c**). Each of the two enrichment conditions was analyzed in triplicates, and the replicates showed high correlations (r = for MECP2 wt and r ≥ 0.99 for MECP2 HM), demonstrating the robustness of each condition. As expected, the input libraries for MECP2 HM and MECP2 wt clustered together, but exhibited lower correlations with the enriched libraries (r ≈ 0.84 - 0.89). Additionally, heatmaps and average profiles showed sharp, reproducible enrichment of MECP2 wt and MECP2 HM at peak centers relative to corresponding input (**Fig. S8**). Collectively, these results confirm that MECP2 enrichment is robust, reproducible across replicates, and clearly distinguishes the different experimental conditions. Peak calling identified 91,481 peaks for MECP2 wt and 149,656 peaks for MECP2 HM (present in at least 2 of 3 technical replicates). Constructing a consensus peak universe revealed that 24% of peaks were unique to MECP2 wt, 54% were unique to MECP2 HM, and 22% were shared between the two conditions (**Fig. 3d**). We used GAT to assess peak overlap with genomic features relative to background (workspace).

Both MECP2 wt- and MECP2 HM-associated peaks were significantly enriched in 3′-untranslated regions (UTRs), coding sequences (CDS), and transcription termination sites

(TTS, ±1 kb). Notably, MECP2 HM peaks showed a significant enrichment for CpG islands compared to MECP2 wt. In contrast, both MECP2 wt and MECP2 HM peaks were significantly depleted (q < 0.05) in rRNA and tRNA genes, as well as in 5′-UTRs and transcription start sites (TSS, ±1 kb) of protein-coding genes (**Fig. S10**).

To put HM-DyadCap into context with existing enrichment methods, we next studied anti-mC and anti-hmC antibodies, being the enrichment probes in MeDIP and hMeDIP protocols. We were first interested in assessing the dyad selectivities of anti-hmC antibodies and their potential use for dyad-specific enrichments. Whereas antibodies for DNA modifications typically require denaturation of the sample DNA (leading to loss of dyad information), some antibodies offer applications with native dsDNA according to the manufacturers. We first tested three different monoclonal antibodies in EMSA with oligonucleotides containing one hmCpG site in single-stranded form, or hybridized to a complement strand to afford a dsDNA with a single hmC/mC dyad. Whereas all antibodies bound the ssDNA (**Fig S9**), none of them showed binding to the dsDNA target even at high concentrations (**Fig. 4a**). However, we identified a polyclonal antibody that was able to bind this dsDNA target at high concentrations, and employed it in EMSA with dsDNA targets containing either an hmC/C, hmC/mC, hmC/hmC or mC/mC CpG dyad. We observed higher affinity for hmC/hmC and hmC/mC dsDNA and only very low or no affinity for hmC/C and mC/mC targets, indicating that this antibody is not suited for selective enrichment of individual hmC dyads (**Fig 4b-c**).

**Figure 4.**
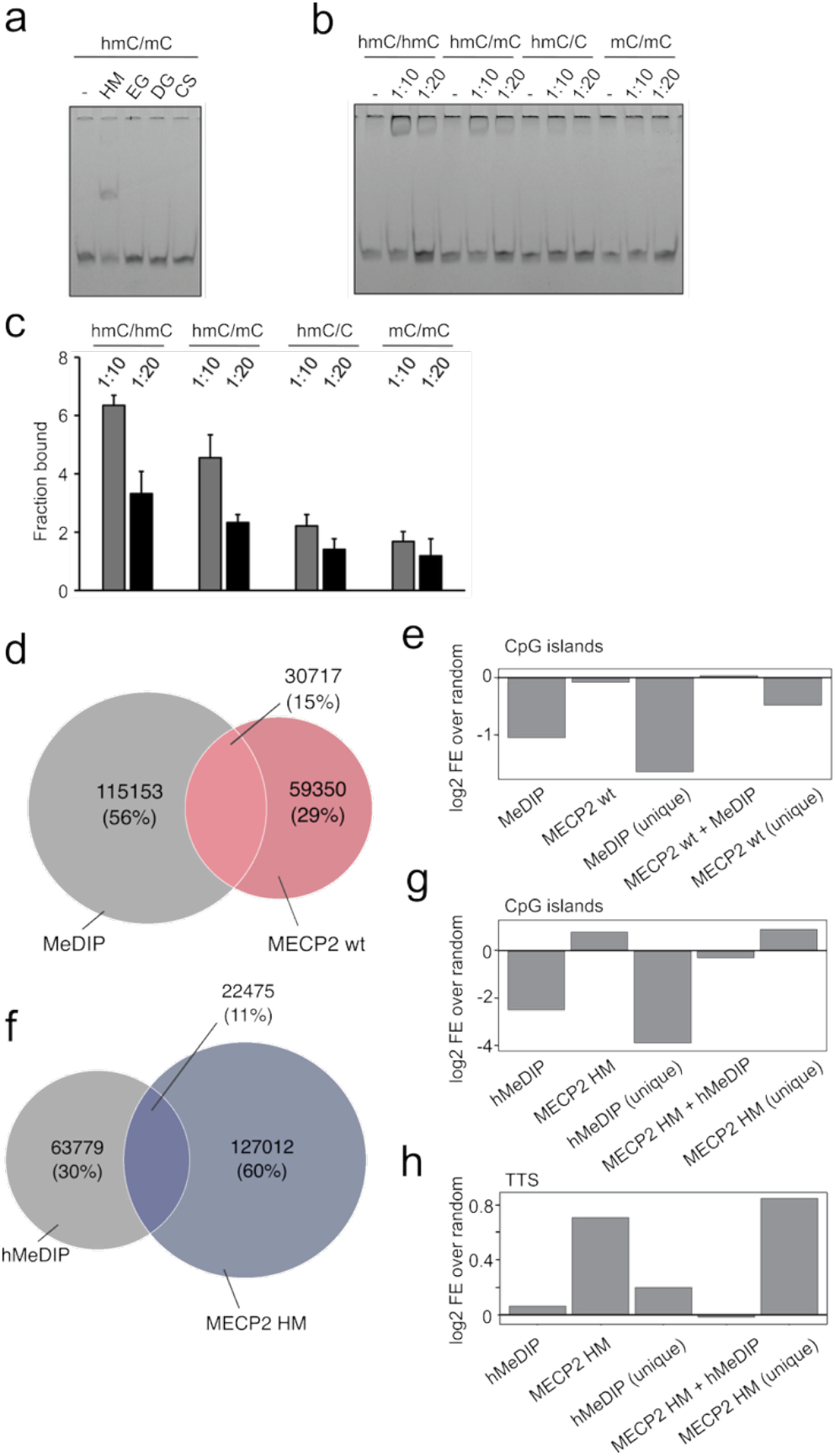
Comparison of MECP2 wt and HM maps with MeDIP and hMeDIP maps. a) EMSA analysis of monoclonal anti-hmC antibodies (1:10 dilution) binding to dsDNA oligos (0.75 pM) bearing a single hmC/mC dyad (for data with hmC-modified ssDNA, see **Fig. S9**). HM: MECP2 HM positive control, EG, DG, CS: antibody supplier (see methods). b) EMSA analysis of polyclonal anti-hmC antibody binding to dsDNA oligo (0.75 pM) containing the indicated CpG dyad modifications. Antibody dilutions on top. c) Bar diagram of quantification of EMSA data from **Fig. 4b**, error bars show standard deviation (n=2). d) Venn diagram for peaks from MeDIP and MECP2 wt enrichments. e) Enrichment/depletion of CpG islands for selected peak sets. f) Venn diagram for peaks from hMeDIP and MECP2 HM enrichments. g-h) Enrichment/depletion of CpG islands and TTS for selected peak sets.

To provide antibody-based reference maps for HM-DyadCap, and to facilitate comparison between the two approaches, we next conducted standard MeDIP and hMeDIP enrichments using denatured, single stranded gDNA of the same mESC batches as used above. After sequencing and mapping, we corrected the dataset with an IgG-only control enrichment according to Lentini et al.^54^. We then assessed the genome-wide overlap between regions enriched with MeDIP and MECP2 wt, respectively. This analysis revealed that 35% of MECP2 wt peaks overlapped with MeDIP. Consistent with this, peak comparison showed that the majority of peaks were unique to MeDIP (56%), whereas MECP2 wt contributed a smaller unique fraction (29%). A total of 30,717 peaks (15%) were shared between the two datasets (**Fig. 4d**), indicating that MECP2 wt covers a subset of genomic regions that is to a good part distinct from MeDIP. Although DIP-seq methods are important tools for mapping modified DNA bases, they have limitations that may account for the discrepancy of the two datasets. These include bias toward low-CG regions^61^, overrepresentation of highly modified regions^62^, nonspecific enrichment of short tandem repeats^54^, and (unlike MECP2 wt), an inability to discriminate between mC/mC and mC/C CpGs. A re-evaluation of DIP-seq approaches showed that 50–99% of enriched regions in datasets without correction by an IgG-only control were false positives, regardless of modification type, cell type, or organism^54^.

To next investigate the genomic distribution of the peak sets of the MeDIP and MECP2 wt experiments, we assessed their enrichment across diverse genomic features. Both datasets showed significant enrichment at CDS, 3′-UTRs, and TTS, while being consistently depleted at rRNA, tRNA genes, 5′-UTRs, and TSS (**Fig. S11**). Notably, the major discrepancy was observed at CpG islands. To better study this discrepancy, we examined enrichment of selected peak subsets (unique to MeDIP, unique to MECP2 wt, and common peaks for both treatments). MeDIP peaks were markedly depleted at CpG islands, whereas MECP2 wt peaks were not, with the former being more pronounced for peaks that were unique to MeDIP (**Fig. 4e**, such a differential enrichment of regions with high versus low CpG densities has previously been observed for the two methods^61,63,64^). Next, comparison of distributions of hMeDIP and MECP2 HM peaks showed that 15% of MECP2 HM peaks overlapped with the hMeDIP peaks. Peak overlap analysis revealed that a high number of regions was unique to MECP2_HM (60%), while hMeDIP contributed a smaller fraction (30%, **Fig. 4f**). A total of 22,475 peaks were shared between the two datasets, suggesting that also MECP2 HM covers a predominantly distinct subset of genomic regions compared to hMeDIP. Notably, CpG islands displayed the most pronounced global difference: MECP2 HM peaks were slightly enriched, whereas hMeDIP peaks were strongly depleted. Importantly, these trends were mainly driven by the peaks that are specific to each treatment (hMeDIP and MECP2 HM), whereas peaks that are common for both treatments showed an intermediary state (**Fig. 4g**). Furthermore, at TTS, MECP2 HM specific peaks were significantly enriched, whereas hMeDIP-specific peaks showed only a slight enrichment (**Fig. 4h**). Collectively, these analyses indicate that MECP2 HM binding may be more selectively enriched at regulatory regions – including CpG islands, 3′-UTRs, and TTS – whereas hMeDIP captures a distinct genomic subsets that may be functionally less specific (**Fig. S11**). Again, whereas MECP2 HM shows selectivity for hmC/mC over other hmC-modified dyads (**Fig. 2a**), hMeDIP is conducted with denatured DNA and expected to enrich hmC regardless of dyad symmetry.

MECP2 HM binds its target hmC/mC dyad with high selectivity over the off-target dyads mC/C, hmC/C, and hmC/hmC (>100-, >125- and ~50-fold respectively; **Fig. 2a**)^33^ that furthermore occur at low levels in mESC and other characterized genomes compared to mC/mC^20,23,25^. MECP2 HM also shows selectivity over mC/mC (10-fold, **Fig. 2a**^33^), but this off-target dyad is particularly abundant^20,23,25^. We introduced an additional control to assess the hmC/mC selectivity of MECP2 HM across the genome. In EMSA experiments, we found that the glucosylation of hmC to glucosyl-hmC (ghmC, **Fig. 5a** and **S1**) by T4 BGT effectively blocks the binding of MECP2 HM (**Fig. 5b** and **S12**).

**Figure 5.**
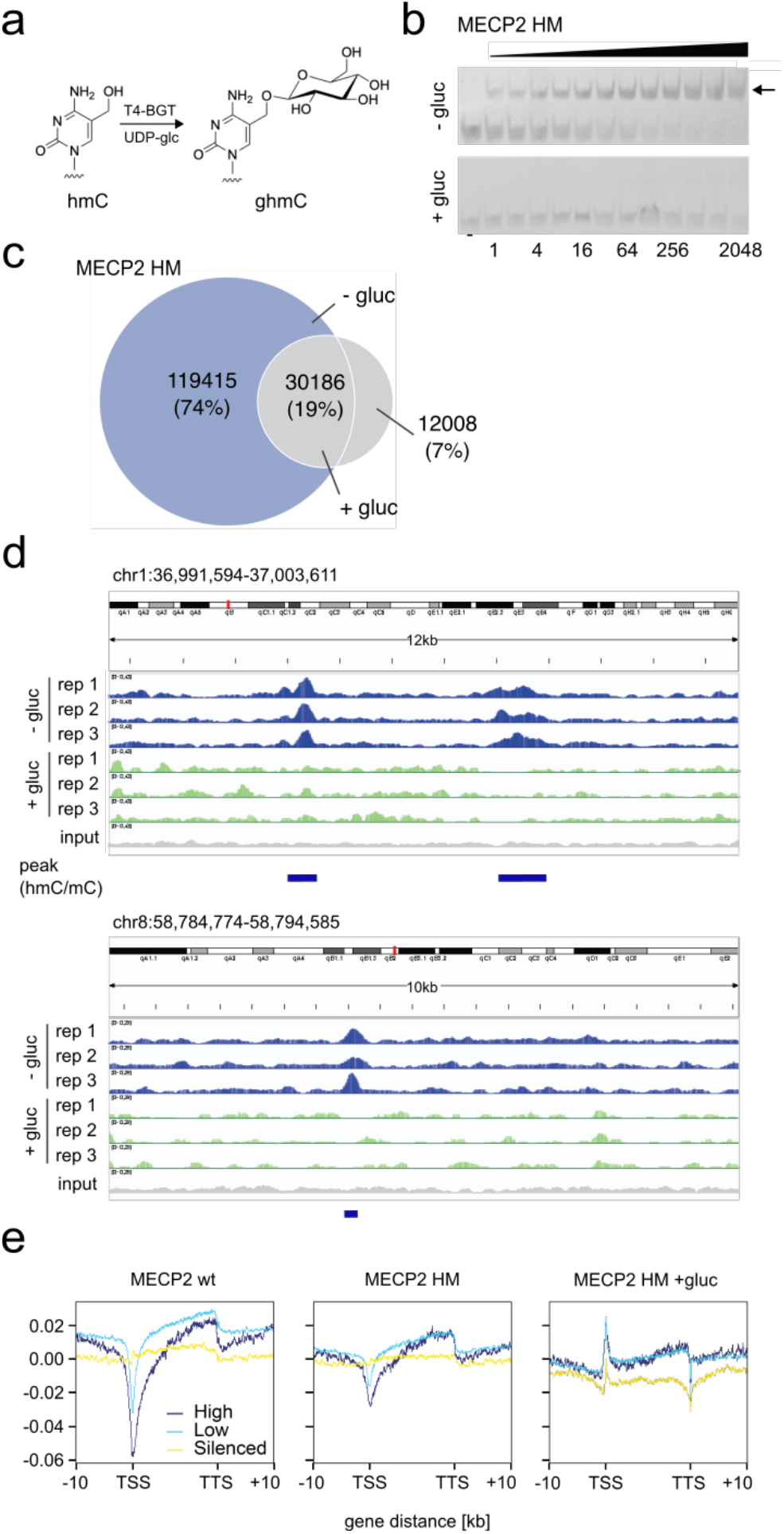
Glucosylation control indicates selective hmC/mC enrichment by MECP2 HM. a) Reaction scheme for T4 BGT-catalyzed glucosylation of hmC. b) EMSA of MECP2 HM binding to dsDNA containing a single hmC/mC dyad and glucosylated or not glucosylated with T4 BGT. c) Venn diagram for peaks obtained from MECP2 HM enrichments with glucosylated or non-glucosyated mESC gDNA. d) Example regions of enrichment signal tracks for non-glucosylated and glucosylated gDNA (shown are three technical replicates each). e) Relative positioning of enrichment signal strengths in genes (+/-10 kb with respect to TSS and TTS) with high, medium or low expression.

Since MECP2 HM shows only very low binding to hmC off-target dyads and mC/mC cannot be glucosylated, enrichment differences between glucosylated and non-glucosylated gDNA should indicate hmC/mC dyads with high confidence. Therefore, we glucosylated part of the same mESC gDNA batch used in prior experiments, and performed MECP2 HM enrichment to verify if peak loss is indeed observed. A comparative peak analysis of these experiments is summarized in the Venn diagram of **Fig. 5c**. Indeed, we observed a significant loss in identified peaks upon glucosylation: we identified only 42,194 peaks (+gluc) vs. 149,601 in the non-glucosylated enrichment (-gluc), indicating that the large majority of peaks enriched by MECP2 HM indeed seems to originate from the selective enrichment of hmC/mC dyads. However, there was also some overlap between both treatments (19%; 30,186 peaks), presumably due to residual background enrichment of clustered off-targets (**Fig. 5c**). **Fig 5d** shows integrative genomics viewer (IGV) signal tracks for MECP2 HM enrichments with glucosylated and non-glucosylated mESC. The data show consistent peak enrichment across technical replicates in both cases, with both overlapping and condition-specific regions (**Fig. S14**). We thereby observed a striking loss of MECP2 HM signals at different loci for the glucosylated control, in agreement with the blocking of MECP2 HM by ghmC observed before (**Fig. 5b**).

The enrichment of hmC within gene bodies has been linked to transcriptional activity in diverse tissues^5,8,9^. To study the relative distribution of enrichment within genes on a global scale, we generated metagene read density profiles aligned to protein-coding genes, clustered according to their expression levels (**Fig 5e**). Profiles were highly consistent across replicates, demonstrating reproducibility of the enrichment patterns (**Fig. S13**). Both MECP2 wt and HM showed robust enrichment across bodies of active compared to silenced genes, with pronounced depletion at TSS and mild depletion at TTS. Enrichment also extended into upstream and downstream flanking regions, consistent with prior reports describing enrichment of hmC^30^ and hmC/mC^23^ around gene bodies of highly expressed genes. In contrast, glucosylation of hmC markedly reduced MECP2 HM enrichment. Density plots and heatmaps revealed a substantial loss of signal across gene bodies, with increased occupancy near TSS and the depletion at TTS. Together, these results indicate that MECP2 wt and HM associate with gene bodies and adjacent regulatory regions, but that the interaction of MECP2 HM is significantly affected by hmC-glucosylation, in agreement with our in vitro EMSA data (**Fig. 5b**).

Enhancers are regulatory elements that control the transcription of distal target genes and play essential roles in cell differentiation by establishing cell type–specific transcriptional programs^65^. TET-mediated oxidation of mC has been shown to modulate enhancer activity during early stages of differentiation^66^, and enhancers consistently exhibit significant levels of hmC across diverse cell types^65^. To assess how the binding of MECP2 variants is influenced by enhancer state, we analyzed the genome-wide distribution of MECP2 variants across enhancer classes defined by Cruz-Molina et. al.^67^. We examined MECP2 occupancy at four enhancer categories: active enhancers, primed enhancers, poised enhancers, and poised-to-active (PoiAct) enhancers (**Fig. S15**). Distinct binding profiles were observed across enhancer states. In general, MECP2 wt and MECP2 HM showed depletion at the center of enhancers, while the MECP2 HM +gluc control showed enrichment. The strength of this effect varied between enhancer types and showed the most pronounced difference between MECP2 wt and HM for active and PoiAct enhancers, where MECP2 HM exhibited a lower relative depletion at the enhancer center than MECP2 wt. Input controls remained flat across all enhancer classes, confirming signal specificity. Overall, these results indicate that MECP2 binding is not uniform but is strongly shaped by enhancer state and MECP2 variant.

## CONCLUSION

In this study, we report HM-DyadCap, a robust method for selectively capturing DNA fragments modified with asymmetric hmC/mC CpG dyads that are frequent marks in mESC and mouse brain tissue. Central to this approach is MECP2 HM, an engineered methyl-CpG-binding domain for that EMSA studies reveal high target affinity and specificity with respect to off-target dinucleotides. We demonstrate that HM-DyadCap enables simple mapping of hmC/mC dyads in a mammalian genome, thus representing the first dyad-specific hmC enrichment method. Comparative mapping studies with established protocols such as wild-type MECP2-based MethylCap as well as MeDIP and hMeDIP reveal differences in the capability of enriching particular genomic features, such as a high enrichment of CpG islands for MECP2 HM as compared to the hMeDIP protocol. The use of glucosylation controls further supports the selectivity of MECP2 HM for hmC/mC dyads. Our mapping experiments reveal that these dyads are enriched in gene bodies and depleted at transcription start sites of protein coding genes in mESC genomes, and that they correlate with active transcription. Moreover, enhancer profiling highlights a dynamic relationship between MECP2 binding and enhancer state, implicating hmC/mC as a potentially important mark in enhancer regulation.

HM-DyadCap may serve as a valuable tool for studying hmC/mC’s regulatory functions and dynamics in development and disease. The ability of dyad-specific enrichment is particularly relevant for generating maps in genomes that exhibit low hmC levels, and for large-scale studies for that whole genome deep sequencing may be cost-prohibitive. HM-DyadCap will therefore be useful for cancer biomarker discovery via effective lower resolution mapping in cancer genomes that have generally low hmC levels. However, the method may also be combined with conversion-based sequencing methods to enable high-resolution maps without the need for whole genome deep sequencing. The MBD of MECP2 can be reengineered to recognize dyads other than hmC/mC^50^, promising an extension to other dyad marks with high relevance in chromatin regulation and cancer development.

## DATA AVAILABILITY

The processed RNA sequencing data reported in this paper was obtained from ENCODE repository under the accession codes: ENCFF827OZU and ENCFF898TDI. The sequencing datasets underlying this study are publicly accessible through ArrayExpress under the respective accession number: E-MTAB-15857 and E-MTAB-15961.

## SUPPLEMENTARY INFORMATION

Associated content 1: Tables S1-S2, Figures S1-S15 (PDF) This material is available free of charge via the Internet.

## AUTHOR CONTRIBUTIONS

L.E. and D. Sc. conducted wet lab experiments and analyzed data. M.Z., A.S. and S.B. analyzed NGS data and contributed to data interpretation. K.K. conducted EMSAs.

B.B. developed methods. S.T. and J.I. supported the analysis of NGS data. C.S. conducted cell culture of mESCs and provided consulting. D. Su., S.B., M.Z., L.E. and K.K. wrote the manuscript with input from all other authors. D. Su. And S.B. supervised the study.

## ACKNOWLEDGEMENT

We thank the Faculty of Chemistry and Chemical Biology of the TU Dortmund University and the International Max-Planck Research School for Living Matter for continuous support. We thank the members of the group for support. We thank Agnieszka Zelisko and Anne-Clémence Veillard from Diagenode for providing the hMeDIP antibody as well as for discussions and input regarding DIP experiments.

## FUNDING

Funded by the Deutsche Forschungsgemeinschaft (SU726/10-1), the Max-Planck Society, the CANTAR program “Netzwerke 2021” and the NRW returning scholars program, both being initiatives of the Ministry of Culture and Science of the State of Northrhine Westphalia. Funded by the European Union (ERC, 101100794 COMBICODE). Views and opinions expressed are those of the author(s) only and do not necessarily reflect those of the European Union or the European Research Council. Neither the European Union nor the granting authority can be held responsible for them.

## CONFLICT OF INTEREST

The authors declare the following competing financial interest: TU Dortmund University has filed a patent application for the engineered MBD employed in the present study (PCT/EP2020/087979, pending). No further competing financial interests have been declared.

## Notes

### Summary of Updates

statement for equal contribution of first two authors has been added

