## Supplementary information for "HM-DyadCap – Capture and Mapping of 5-Hydroxymethylcytosine/5-Methylcytosine CpG Dyads in Mammalian DNA"

[c]Genomic Services, Qiagen, Manchester (United Kingdom).

This PDF file includes:

**SI Table 1-2**

**SI Fig. 1-15**

**Table S1.** Oligonucleotides used in this study.

| Name | Description | Sequence 5’ -> 3’ |
|---|---|---|
| o4123 | qPCR, 5‘-Phos | TCAGCCTTTCATTGATTGCG |
| o4124 | qPCR | CTTCTCCTTTACTAGTGAATTC |
| o4368 | qPCR | CTCTTCTGCCTGCTGACCTTTG |
| o4371 | qPCR | CTTTCATTGATTGCGGATTCCAGAATTCACTAGTAAAGGAGAAGTTGGCTACAGCAA |
| o4372 | qPCR | ACCACCCTGTTGCTGTAGCCAA |
| o4373 | qPCR | CTTTCATTGATTGCGCACGATAGAATTCACTAGTAAAGGAGAAGGAACCGCTCATTG |
| o4374 | qPCR | CACCATTGGCAATGAGCGGTTC |
| o4392 | qPCR | CTTTCATTGATTGCGTCACAGAGAATTCACTAGTAAAGGAGAAGTGTAGCCCTCTGT |
| o4393 | qPCR | CTTGAGCACACAGAGGGCTACA |
| o4538 | NGS adapter | ACACTCTTTCCCTACACGACGCTCTTCCGATCTCTTCCTGGCACGAGTCACCCTTTCATTC ATTCCC |
| o4539 | NGS adapter | CCACGAGATAAGAGGATGGCAAACAGCTATGACNNNNNNNNNNCTTCTCCTTTACTACTCA ATTC |
| o4627 | template for o4372 | CTCTTCTGCCTGCTGACCTTTGTCAGCCTTTCATTGATTGCGGATTCCAGAATTCACTAGT AAAGGAGAAGTTGGCTACAGCAA |
| o4628 | template for o4374 | CTCTTCTGCCTGCTGACCTTTGTCAGCCTTTCATTGATTGCGGATTCCAGAATTCACTAGT AAAGGAGAAGTTGGCTACAGCAA |
| o4629 | template for o4393 | CTCTTCTGCCTGCTGACCTTTGTCAGCCTTTCATTGATTGCGTCACAGAGAATTCACTAGT AAAGGAGAAGTGTAGCCCTCTGT |
| o4675 | probe; N,N,N,N=5hmC; 5'-FAM | TCTTCNGTTTCCTCAGCNGAAGGCTCGAGTCTTCNGTTTCCAAGCTTCAGCNGAAGGCTCT TCTGCCTGCTGACCTTTG |
| o4677 | probe, N,N,N,N=5mC; 5'-FAM | TCTTCNGTTTCCTCAGCNGAAGGCTCGAGTCTTCNGTTTCCAAGCTTCAGCNGAAGGCTCT TCTGCCTGCTGACCTTTG |
| o4727 | probe; N,N,N,N=5mC; 5'-Phos | CAAAGGTCAGCAGGCAGAAGAGCCTTNGGCTGAAGCTTGGAAANGGAAGACTCGAGCCTTN GGCTGAGGAAANGGAAGA |
| o4728 | probe | TCTTCCGTTTCCTCAGCCGAAGGCTCGAGTCTTCCGTTTCCAAGCTTCAGCCGAAGGCTCT TCTGCCTGCTGACCTTTG |
| o4729 | probe | CAAAGGTCAGCAGGCAGAAGAGCCTTCGGCTGAAGCTTGGAAACGGAAGACTCGAGCCTTC GGCTGAGGAAACGGAAGA |
| o5241 | 8NX probe template | TCACCCTTTCATTCATTCCCTCCACACAACAACCATTCTTNNNNCGNNNNAATGTGAGGAG GGTGTTATAGAATTGAGTAGTAAAGGAGAAG |
| o5242 | PCR primer, N=biotin TEG | NTCACCCTTTCATTCATTCCCTCCACACAACAACCATTC |
| o5243 | PCR primer | CTTCTCCTTTACTACTCAATTCTATAACACCCTCCTCAC |
| o2968 | EMSA probe competitor_A | AAAAAAAAAAAAAAAAAAAAAAAA |
| o2969 | EMSA probe competitor_T | TTTTTTTTTTTTTTTTTTTTTTTT |
| o3112 | EMSA probe X=hmC | TTTTTTTTTTTXGTTTTTTTTTTT |
| o3115 | EMSA probe 5'-FAM; X=hmC | AAAAAAAAAAAXGAAAAAAAAAAA |
| o2909 | EMSA probe X=mC | TTTTTTTTTTTXGTTTTTTTTTTT |
| o5990 | EMSA probe 5'-FAM; X=mC | AAAAAAAANNNXANNNAAAAAAAA |
| o5991 | EMSA probe | TTTTTTTTNNNTGNNNTTTTTTTT |

| | | |
|---|---|---|
| o5992 | EMSA probe<br>5'-FAM; X=mC | AAAAAAAANNNXTNNNAAAAAAAA |
| o5993 | EMSA probe | TTTTTTTTNNNAGNNNTTTTTTTT |
| o5994 | EMSA probe<br>5'-FAM; X=mC | AAAAAAAANNNXCNNNAAAAAAAA |
| o5995 | EMSA probe | TTTTTTTTNNNGGNNNTTTTTTTT |
| o5996 | EMSA probe<br>5'-FAM; X=mC | AAAAAAAANNNXGNNNAAAAAAAA |
| o5997 | EMSA probe<br>X=mC | TTTTTTTTNNNXGNNNTTTTTTTT |
| o5998 | EMSA probe<br>5'-FAM | AAAAAAAANNNCTNNNAAAAAAAA |
| o5999 | EMSA probe<br>5'-FAM | AAAAAAAANNNCANNNAAAAAAAA |
| o6000 | EMSA probe<br>5'-FAM | AAAAAAAANNNCCNNNAAAAAAAA |
| o6001 | EMSA probe<br>5'-FAM | AAAAAAAANNNCGNNNAAAAAAAA |
| o6002 | EMSA probe | TTTTTTTTNNNCGNNNTTTTTTTT |
| o6003 | EMSA probe<br>5'-FAM | AAAAAAAANNNAANNNAAAAAAAA |
| o6004 | EMSA probe | TTTTTTTTNNNTTNNNTTTTTTTT |
| o6005 | EMSA probe<br>5'-FAM | AAAAAAAANNNATNNNAAAAAAAA |
| o6006 | EMSA probe | TTTTTTTTNNNATNNNTTTTTTTT |
| o6007 | EMSA probe<br>5'-FAM | AAAAAAAANNNACNNNAAAAAAAA |
| o6008 | EMSA probe | TTTTTTTTNNNGTNNNTTTTTTTT |
| o6009 | EMSA probe<br>5'-FAM | AAAAAAAANNNTANNNAAAAAAAA |
| o6010 | EMSA probe | TTTTTTTTNNNTANNNTTTTTTTT |
| o6011 | EMSA probe<br>5'-FAM | AAAAAAAANNNTCNNNAAAAAAAA |
| o6012 | EMSA probe | TTTTTTTTNNNGANNNTTTTTTTT |
| o6013 | EMSA probe<br>5'-FAM | AAAAAAAANNNGCNNNAAAAAAAA |
| o6014 | EMSA probe | TTTTTTTTNNNGCNNNTTTTTTTT |
| o6411 | EMSA probe<br>5'-FAM; X=mC | AAAAAAAANNNXGNNNAAAAAAAA |
| o5906 | EMSA probe<br>5'-FAM; X=mC | TAGGCCAXGTGGGAGG |
| o5907 | EMSA probe<br>X=hmC | CCTCCCAXGTGGCCTA |
| o5918 | EMSA probe | CCTCCCACGTGGCCTA |
| o5926 | EMSA probe<br>5'-FAM; X=mC | TAGGCCAXGTGGGAGG |
| o5927 | EMSA probe<br>X=mC | CCTCCCAXGTGGCCTA |
| o4277 | EMSA probe<br>5'-FAM | AAAAAAAAAAATGAAAAAAAAAAA |
| o4278 | EMSA probe<br>X=mC | TTTTTTTTTTTXATTTTTTTTTTT |

**Table S2**. Sequencing data of all sequenced libraries.

| Sample ID (File) | Library Prep Kit | Input DNA (ng) | Insert Size (bp) | Read Leng | Total Read | Raw Bases | % Q30 Bas | GC Conten | Mapping Rate (% | Duplication Rate (% |
|---|---|---|---|---|---|---|---|---|---|---|
| ghmC_TAYN_CAP_1_MKDL240002982-1A_L6_L2_1.fq.gz | NEBNext Ultra II DN/ | 250 | 200 | 150 | 64282894 | 9642434100 | 93,23 | 43,87 | | |
| ghmC_TAYN_CAP_1_MKDL240002982-1A_L6_L2_2.fq.gz | NEBNext Ultra II DN/ | 250 | 200 | 150 | 64282894 | 9642434100 | 91,86 | 44,14 | 95,04 | 28,82177843 |
| ghmC_TAYN_CAP_2_MKDL240002982-1A_L6_L2_1.fq.gz | NEBNext Ultra II DN/ | 250 | 200 | 150 | 64896802 | 9734520300 | 93,22 | 43,88 | | |
| ghmC_TAYN_CAP_2_MKDL240002982-1A_L6_L2_2.fq.gz | NEBNext Ultra II DN/ | 250 | 200 | 150 | 64896802 | 9734520300 | 91,79 | 43,96 | 95,14 | 29,2052956 |
| ghmC_TAYN_CAP_3_MKDL240002982-1A_L6_L2_1.fq.gz | NEBNext Ultra II DN/ | 250 | 200 | 150 | 63199407 | 9479911050 | 93,65 | 43,86 | | |
| ghmC_TAYN_CAP_3_MKDL240002982-1A_L6_L2_2.fq.gz | NEBNext Ultra II DN/ | 250 | 200 | 150 | 63199407 | 9479911050 | 91,95 | 43,75 | 95,23 | 31,58068788 |
| Methyl-CAP-1_R_1.fq.gz | NEBNext Ultra II DN/ | 250 | 200 | 150 | 127020593 | 19053088950 | 93,4 | 42,58 | | |
| Methyl-CAP-1_R_2.fq.gz | NEBNext Ultra II DN/ | 250 | 200 | 150 | 127020593 | 19053088950 | 90,8 | 42,6 | 96,28 | 24,71133967 |
| Methyl-CAP-2_R_1.fq.gz | NEBNext Ultra II DN/ | 250 | 200 | 150 | 157923080 | 23688462000 | 93,28 | 42,6 | | |
| Methyl-CAP-2_R_2.fq.gz | NEBNext Ultra II DN/ | 250 | 200 | 150 | 157923080 | 23688462000 | 91,82 | 42,58 | 96,3 | 25,75834775 |
| Methyl-CAP-3_R_1.fq.gz | NEBNext Ultra II DN/ | 250 | 200 | 150 | 138393530 | 20759029500 | 93,49 | 42,6 | | |
| Methyl-CAP-3_R_2.fq.gz | NEBNext Ultra II DN/ | 250 | 200 | 150 | 138393530 | 20759029500 | 92,25 | 42,61 | 96,38 | 25,48510666 |
| TAYN-CAP-1_R_1.fq.gz | NEBNext Ultra II DN/ | 250 | 200 | 150 | 184981617 | 27747242550 | 93,4 | 41,32 | | |
| TAYN-CAP-1_R_2.fq.gz | NEBNext Ultra II DN/ | 250 | 200 | 150 | 184981617 | 27747242550 | 92,04 | 41,32 | 96,59 | 33,75818779 |
| TAYN-Cap-2_R_1.fq.gz | NEBNext Ultra II DN/ | 250 | 200 | 150 | 147450685 | 22117602750 | 93,27 | 41,31 | | |
| TAYN-Cap-2_R_2.fq.gz | NEBNext Ultra II DN/ | 250 | 200 | 150 | 147450685 | 22117602750 | 92,57 | 41,32 | 96,63 | 32,19847944 |
| TAYN-Cap-3_R_1.fq.gz | NEBNext Ultra II DN/ | 250 | 200 | 150 | 144005396 | 21600809400 | 93,17 | 41,28 | | |
| TAYN-Cap-3_R_2.fq.gz | NEBNext Ultra II DN/ | 250 | 200 | 150 | 144005396 | 21600809400 | 91,94 | 41,3 | 96,56 | 31,78881241 |
| TAYN-input_R_1.fq.gz | NEBNext Ultra II DN/ | 250 | 200 | 150 | 154237980 | 23135697000 | 93,39 | 41,88 | | |
| TAYN-input_R_2.fq.gz | NEBNext Ultra II DN/ | 250 | 200 | 150 | 154237980 | 23135697000 | 92,78 | 41,88 | 96,65 | 18,67804527 |
| wt-input_R_1.fq.gz | NEBNext Ultra II DN/ | 250 | 200 | 150 | 133293660 | 19994049000 | 93,22 | 41,86 | | |
| wt-input_R_2.fq.gz | NEBNext Ultra II DN/ | 250 | 200 | 150 | 133293660 | 19994049000 | 92,09 | 41,84 | 96,65 | 17,01109214 |
| R3_WT3_merge_2.fq.gz | NEBNext Ultra II DN/ | 250 | 209 | 150 | 47383768 | 7107565200 | 86,45 | 43,42 | | |
| R3_WT3_merge_1.fq.gz | NEBNext Ultra II DN/ | 250 | 209 | 150 | 47383768 | 7107565200 | 88,11 | 44,59 | 60,88 | 30,37651427 |
| R3_WT2_merge_2.fq.gz | NEBNext Ultra II DN/ | 250 | 209 | 150 | 52389202 | 7858380300 | 86,59 | 42,98 | | |
| R3_WT2_merge_1.fq.gz | NEBNext Ultra II DN/ | 250 | 209 | 150 | 52389202 | 7858380300 | 87,7 | 43,63 | 62,12 | 30,73286866 |
| R3_WT1_merge_2.fq.gz | NEBNext Ultra II DN/ | 250 | 209 | 150 | 53868099 | 8080214850 | 87,56 | 42,99 | | |
| R3_WT1_merge_1.fq.gz | NEBNext Ultra II DN/ | 250 | 209 | 150 | 53868099 | 8080214850 | 88,65 | 44,06 | 61,66 | 29,74790317 |
| R3_TAYNgluc3_merge_2.fq.gz | NEBNext Ultra II DN/ | 250 | 285 | 150 | 82530386 | 12379557900 | 91,6 | 42,17 | | |
| R3_TAYNgluc3_merge_1.fq.gz | NEBNext Ultra II DN/ | 250 | 285 | 150 | 82530386 | 12379557900 | 92,71 | 42,13 | 84,66 | 35,38286924 |
| R3_TAYNgluc2_merge_2.fq.gz | NEBNext Ultra II DN/ | 250 | 285 | 150 | 85605070 | 12840760500 | 91,94 | 41,96 | | |
| R3_TAYNgluc2_merge_1.fq.gz | NEBNext Ultra II DN/ | 250 | 285 | 150 | 85605070 | 12840760500 | 92,74 | 41,89 | 84,58 | 35,38944914 |
| R3_TAYNgluc1_merge_2.fq.gz | NEBNext Ultra II DN/ | 250 | 285 | 150 | 76613797 | 11492069550 | 92,23 | 41,8 | | |
| R3_TAYNgluc1_merge_1.fq.gz | NEBNext Ultra II DN/ | 250 | 285 | 150 | 76613797 | 11492069550 | 93,14 | 41,73 | 84,7 | 37,27373069 |
| R3_TAYN3_merge_2.fq.gz | NEBNext Ultra II DN/ | 250 | 209 | 150 | 45835560 | 6875334000 | 84,93 | 43,11 | | |
| R3_TAYN3_merge_1.fq.gz | NEBNext Ultra II DN/ | 250 | 209 | 150 | 45835560 | 6875334000 | 86,63 | 44,96 | 59,48 | 23,90566032 |
| R3_TAYN2_merge_2.fq.gz | NEBNext Ultra II DN/ | 250 | 209 | 150 | 51778983 | 7766847450 | 85,1 | 43,04 | | |
| R3_TAYN2_merge_1.fq.gz | NEBNext Ultra II DN/ | 250 | 209 | 150 | 51778983 | 7766847450 | 86,5 | 44,47 | 60,59 | 24,26766815 |
| R3_TAYN1_merge_2.fq.gz | NEBNext Ultra II DN/ | 250 | 209 | 150 | 51323295 | 7698494250 | 86,65 | 42,24 | | |
| R3_TAYN1_merge_1.fq.gz | NEBNext Ultra II DN/ | 250 | 209 | 150 | 51323295 | 7698494250 | 87,18 | 42,7 | 62,4 | 24,57100711 |
| R3_input_unmodified_merge_2.fq.gz | NEBNext Ultra II DN/ | 250 | 209 | 150 | 46833400 | 7025010000 | 87,06 | 42,25 | | |
| R3_input_unmodified_merge_1.fq.gz | NEBNext Ultra II DN/ | 250 | 209 | 150 | 46833400 | 7025010000 | 87,33 | 43,18 | 61,42 | 19,14342731 |
| R3_input_gluc_merge_2.fq.gz | NEBNext Ultra II DN/ | 250 | 285 | 150 | 83586786 | 12538017900 | 91,71 | 42,37 | | |
| R3_input_gluc_merge_1.fq.gz | NEBNext Ultra II DN/ | 250 | 285 | 150 | 83586786 | 12538017900 | 92,47 | 42,24 | 83,83 | 28,00168322 |
| R2_WT3_merge_2.fq.gz | NEBNext Ultra II DN/ | 250 | 223 | 150 | 48608378 | 7291256700 | 88,68 | 43,82 | | |
| R2_WT3_merge_1.fq.gz | NEBNext Ultra II DN/ | 250 | 223 | 150 | 48608378 | 7291256700 | 91,3 | 44,54 | 76,71 | 24,1235632 |
| R2_WT2_merge_2.fq.gz | NEBNext Ultra II DN/ | 250 | 223 | 150 | 56747427 | 8512114050 | 87,7 | 43,61 | | |
| R2_WT2_merge_1.fq.gz | NEBNext Ultra II DN/ | 250 | 223 | 150 | 56747427 | 8512114050 | 90,09 | 44,14 | 76,66 | 25,18976544 |
| R2_WT1_merge_2.fq.gz | NEBNext Ultra II DN/ | 250 | 223 | 150 | 53913633 | 8087044950 | 89,03 | 43,55 | | |
| R2_WT1_merge_1.fq.gz | NEBNext Ultra II DN/ | 250 | 223 | 150 | 53913633 | 8087044950 | 91,27 | 44,23 | 77,39 | 24,58028754 |
| R2_TAYNgluc3_merge_2.fq.gz | NEBNext Ultra II DN/ | 250 | 277 | 150 | 63527993 | 9529198950 | 91,39 | 42,49 | | |
| R2_TAYNgluc3_merge_1.fq.gz | NEBNext Ultra II DN/ | 250 | 277 | 150 | 63527993 | 9529198950 | 92,94 | 42,52 | 88,54 | 30,72548556 |
| R2_TAYNgluc2_merge_2.fq.gz | NEBNext Ultra II DN/ | 250 | 277 | 150 | 72174316 | 10826147400 | 90,97 | 42,27 | | |
| R2_TAYNgluc2_merge_1.fq.gz | NEBNext Ultra II DN/ | 250 | 277 | 150 | 72174316 | 10826147400 | 92,45 | 42,3 | 88,4 | 34,74521282 |
| R2_TAYNgluc1_merge_2.fq.gz | NEBNext Ultra II DN/ | 250 | 277 | 150 | 71073537 | 10661030550 | 91,43 | 42,55 | | |
| R2_TAYNgluc1_merge_1.fq.gz | NEBNext Ultra II DN/ | 250 | 277 | 150 | 71073537 | 10661030550 | 93,01 | 42,67 | 88,41 | 32,22490836 |
| R2_TAYN3_merge_2.fq.gz | NEBNext Ultra II DN/ | 250 | 223 | 150 | 52255680 | 7838352000 | 87,65 | 43,39 | | |
| R2_TAYN3_merge_1.fq.gz | NEBNext Ultra II DN/ | 250 | 223 | 150 | 52255680 | 7838352000 | 90,1 | 44,49 | 74,54 | 22,33024299 |
| R2_TAYN2_merge_2.fq.gz | NEBNext Ultra II DN/ | 250 | 223 | 150 | 49723964 | 7458594600 | 85,92 | 43,33 | | |
| R2_TAYN2_merge_1.fq.gz | NEBNext Ultra II DN/ | 250 | 223 | 150 | 49723964 | 7458594600 | 88,27 | 44,29 | 74,37 | 21,10740738 |
| R2_TAYN1_merge_2.fq.gz | NEBNext Ultra II DN/ | 250 | 223 | 150 | 56928185 | 8539227750 | 87,73 | 43,28 | | |
| R2_TAYN1_merge_1.fq.gz | NEBNext Ultra II DN/ | 250 | 223 | 150 | 56928185 | 8539227750 | 89,9 | 43,85 | 75,92 | 22,97079035 |
| R2_input_unmodified_merge_2.fq.gz | NEBNext Ultra II DN/ | 250 | 223 | 150 | 42387000 | 6358050000 | 85,91 | 43,35 | | |
| R2_input_unmodified_merge_1.fq.gz | NEBNext Ultra II DN/ | 250 | 223 | 150 | 42387000 | 6358050000 | 88,27 | 44,76 | 72,7 | 16,22863373 |
| R2_input_gluc_merge_2.fq.gz | NEBNext Ultra II DN/ | 250 | 277 | 150 | 66584038 | 9987605700 | 89,56 | 43,16 | | |
| R2_input_gluc_merge_1.fq.gz | NEBNext Ultra II DN/ | 250 | 277 | 150 | 66584038 | 9987605700 | 91,28 | 43,42 | 86,34 | 24,82145739 |
| R1_WT3_merge_2.fq.gz | NEBNext Ultra II DN/ | 250 | 200 | 150 | 40575509 | 6086326350 | 86,62 | 43,35 | | |
| R1_WT3_merge_1.fq.gz | NEBNext Ultra II DN/ | 250 | 200 | 150 | 40575509 | 6086326350 | 89,12 | 45,47 | 64,32 | 27,52046698 |
| R1_WT2_merge_2.fq.gz | NEBNext Ultra II DN/ | 250 | 200 | 150 | 53160457 | 7974068550 | 87,66 | 42,83 | | |
| R1_WT2_merge_1.fq.gz | NEBNext Ultra II DN/ | 250 | 200 | 150 | 53160457 | 7974068550 | 89,39 | 43,95 | 66,38 | 33,02907436 |
| R1_WT1_merge_2.fq.gz | NEBNext Ultra II DN/ | 250 | 200 | 150 | 47929501 | 7189425150 | 88,36 | 42,87 | | |
| R1_WT1_merge_1.fq.gz | NEBNext Ultra II DN/ | 250 | 200 | 150 | 47929501 | 7189425150 | 90,51 | 44,82 | 63,94 | 30,94089559 |
| R1_TAYNgluc3_merge_2.fq.gz | NEBNext Ultra II DN/ | 250 | 242 | 150 | 76116333 | 11417449950 | 91,34 | 41,95 | | |
| R1_TAYNgluc3_merge_1.fq.gz | NEBNext Ultra II DN/ | 250 | 242 | 150 | 76116333 | 11417449950 | 92,3 | 42,39 | 83,25 | 35,9286951 |
| R1_TAYNgluc2_merge_2.fq.gz | NEBNext Ultra II DN/ | 250 | 242 | 150 | 63949953 | 9592492950 | 89,72 | 42,47 | | |
| R1_TAYNgluc2_merge_1.fq.gz | NEBNext Ultra II DN/ | 250 | 242 | 150 | 63949953 | 9592492950 | 91,04 | 43,04 | 81,26 | 33,44220422 |
| R1_TAYNgluc1_merge_2.fq.gz | NEBNext Ultra II DN/ | 250 | 242 | 150 | 66893965 | 10034094750 | 91,04 | 41,75 | | |
| R1_TAYNgluc1_merge_1.fq.gz | NEBNext Ultra II DN/ | 250 | 242 | 150 | 66893965 | 10034094750 | 91,99 | 42,08 | 83,19 | 37,06214128 |
| R1_TAYN3_merge_2.fq.gz | NEBNext Ultra II DN/ | 250 | 200 | 150 | 53201776 | 7980266400 | 87,14 | 42,65 | | |
| R1_TAYN3_merge_1.fq.gz | NEBNext Ultra II DN/ | 250 | 200 | 150 | 53201776 | 7980266400 | 89,08 | 44,76 | 61,42 | 24,88855753 |
| R1_TAYN2_merge_2.fq.gz | NEBNext Ultra II DN/ | 250 | 200 | 150 | 40133886 | 6020082900 | 85,98 | 42,78 | | |
| R1_TAYN2_merge_1.fq.gz | NEBNext Ultra II DN/ | 250 | 200 | 150 | 40133886 | 6020082900 | 87,08 | 44,5 | 63,85 | 21,90078638 |
| R1_TAYN1_merge_2.fq.gz | NEBNext Ultra II DN/ | 250 | 200 | 150 | 48528862 | 7279329300 | 86,52 | 42,49 | | |
| R1_TAYN1_merge_1.fq.gz | NEBNext Ultra II DN/ | 250 | 200 | 150 | 48528862 | 7279329300 | 87,7 | 44,05 | 64,96 | 23,60141571 |
| R1_input_unmodified_merge_2.fq.gz | NEBNext Ultra II DN/ | 250 | 200 | 150 | 30128448 | 4519267200 | 84,26 | 42,92 | | |
| R1_input_unmodified_merge_1.fq.gz | NEBNext Ultra II DN/ | 250 | 200 | 150 | 30128448 | 4519267200 | 85,27 | 45,79 | 61,86 | 14,77448046 |
| R1_input_gluc_merge_2.fq.gz | NEBNext Ultra II DN/ | 250 | 242 | 150 | 68911798 | 10336769700 | 90,15 | 42,6 | | |
| R1_input_gluc_merge_1.fq.gz | NEBNext Ultra II DN/ | 250 | 242 | 150 | 68911798 | 10336769700 | 91,24 | 42,94 | 81,26 | 30,70395549 |
| Me-DIP-3_2.fq.gz | MagMeDIP-seq Pack | 1000 | 200 | 150 | 122579688 | 18386953200 | 91,81 | 42,35 | | |
| Me-DIP-3_1.fq.gz | MagMeDIP-seq Pack | 1000 | 200 | 150 | 122579688 | 18386953200 | 92,2 | 42,46 | 94,97 | 37,34636655 |
| Me-DIP-2_2.fq.gz | MagMeDIP-seq Pack | 1000 | 200 | 150 | 96283985 | 14442597750 | 89,45 | 42,35 | | |
| Me-DIP-2_1.fq.gz | MagMeDIP-seq Pack | 1000 | 200 | 150 | 96283985 | 14442597750 | 92,11 | 42,32 | 94,92 | 38,93437539 |
| Me-DIP-1_2.fq.gz | MagMeDIP-seq Pack | 1000 | 200 | 150 | 130872017 | 19630802550 | 89,9 | 42,41 | | |
| Me-DIP-1_1.fq.gz | MagMeDIP-seq Pack | 1000 | 200 | 150 | 130872017 | 19630802550 | 92,31 | 42,51 | 94,8 | 39,47851424 |
| Input-Me-DIP_2.fq.gz | MagMeDIP-seq Pack | 1000 | 200 | 150 | 240112268 | 36016840200 | 92,74 | 42,48 | | |
| Input-Me-DIP_1.fq.gz | MagMeDIP-seq Pack | 1000 | 200 | 150 | 240112268 | 36016840200 | 93,88 | 42,52 | 96,29 | 14,09625648 |
| Input-hMe-DIP_2.fq.gz | MagMeDIP-seq Pack | 1000 | 200 | 150 | 141600132 | 21240019800 | 92,99 | 42,16 | | |
| Input-hMe-DIP_1.fq.gz | MagMeDIP-seq Pack | 1000 | 200 | 150 | 141600132 | 21240019800 | 93,26 | 42,25 | 96,18 | 12,67786965 |
| hMe-DIP-3_2.fq.gz | MagMeDIP-seq Pack | 1000 | 200 | 150 | 106998663 | 16049799450 | 87,87 | 44,21 | | |
| hMe-DIP-3_1.fq.gz | MagMeDIP-seq Pack | 1000 | 200 | 150 | 106998663 | 16049799450 | 88,04 | 44,2 | 92,3 | 20,50171105 |
| hMe-DIP-2_2.fq.gz | MagMeDIP-seq Pack | 1000 | 200 | 150 | 121603198 | 18240479700 | 88,28 | 44,32 | | |
| hMe-DIP-2_1.fq.gz | MagMeDIP-seq Pack | 1000 | 200 | 150 | 121603198 | 18240479700 | 87,94 | 44,42 | 92,35 | 21,61268104 |
| hMe-DIP-1_2.fq.gz | MagMeDIP-seq Pack | 1000 | 200 | 150 | 122621105 | 18393165750 | 87,3 | 43,64 | | |
| hMe-DIP-1_1.fq.gz | MagMeDIP-seq Pack | 1000 | 200 | 150 | 122621105 | 18393165750 | 87,12 | 43,67 | 90,85 | 34,76021755 |

hmC/mC

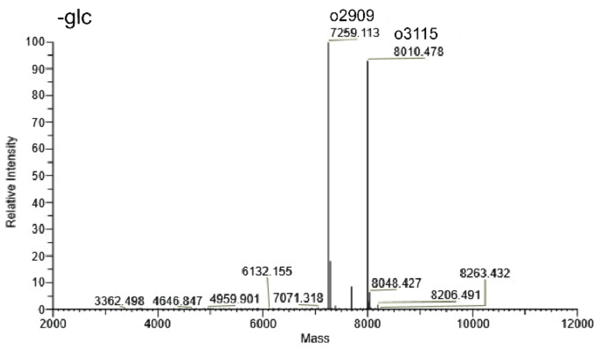


| Sequence Name | Theoretical Mass, Da | Monoisotopic Mass, Da | Mass error, ppm | Relative Abundance, % |
|---|---|---|---|---|
| o2909 | 7259.172 | 7259.113 | 8.0 | 100 |
| o3115 | 8010.540 | 8010.478 | 7.8 | 93.14 |

ghmC/mC

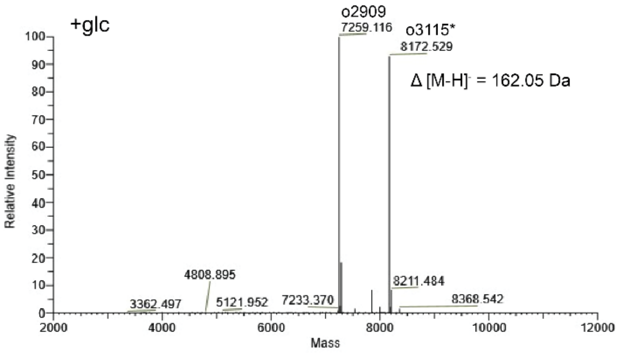


| Sequence Name | Theoretical Mass, Da | Monoisotopic Mass, Da | Mass error, ppm | Relative Abundance, % |
|---|---|---|---|---|
| o2909 | 7259.172 | 7259.116 | 7.7 | 100 |
| o3115 | 8010.540 | NA | NA | NA |
| o3115* | 8172.593 | 8172.529 | 7.8 | 92.93 |

hmC/hmC

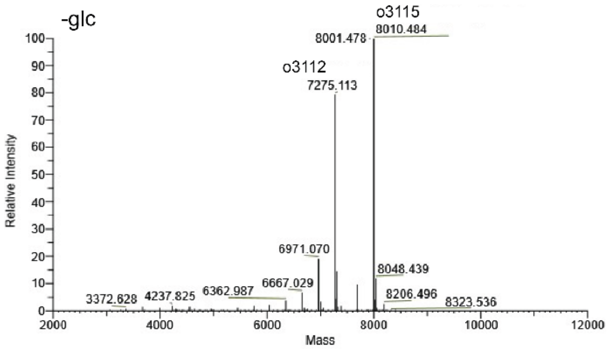


| Sequence Name | Theoretical Mass, Da | Monoisotopic Mass, Da | Mass error, ppm | Relative Abundance, % |
|---|---|---|---|---|
| o3112 | 7275.167 | 7275.113 | 7.3 | 79.56 |
| o3115 | 8010.540 | 8010.484 | 6.9 | 100 |

ghmC/ghmC

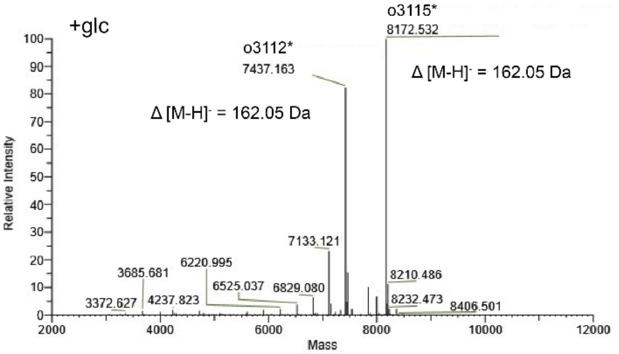


| Sequence Name | Theoretical Mass, Da | Monoisotopic Mass, Da | Mass error, ppm | Relative Abundance, % |
|---|---|---|---|---|
| o3112* | 7437.219 | 7437.163 | 7.6 | 82.65 |
| o3112 | 7275.167 | NA | NA | NA |
| o3115* | 8172.593 | 8172.532 | 7.4 | 100 |
| o3115 | 8010.540 | NA | NA | NA |

**Fig. S1**. T4 β-glucosyltransferase-mediated 5-hydroxymethylcytosine modification analysis using LC-MS. Deconvoluted mass spectra of hmC/mC and hmC/hmC CpG dyad containing 24-mer DNA duplexes before (left) and after (right) glucosylation.

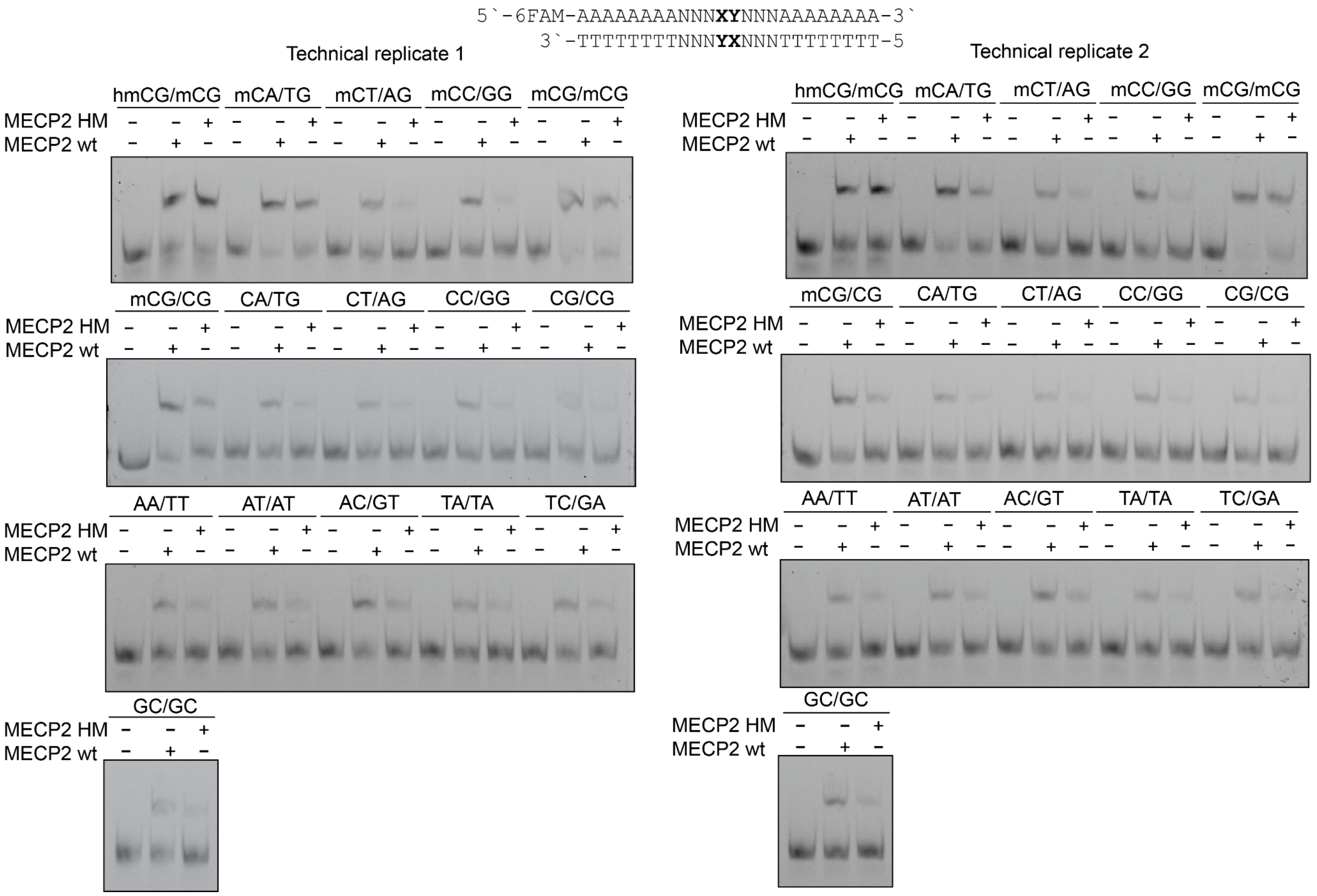


**Fig. S2**. Two technical replicates of electrophoretic mobility shift (EMSA) gel images showing the binding of 100 nM recombinantly expressed MECP2 HM and MECP2 wt proteins to 2 nM of 24-mer DNA duplexes containing a random sequence upstream and downstream of the indicated dinucleotides.

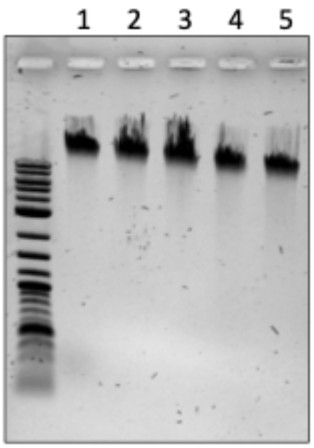


**Fig. S3**. Agarose gel analysis of gDNA isolated from mESC cells (E14TG2A)

| Rep | input gDNA | Sheared gDNA | yield | Avg. fragment length |
|---|---|---|---|---|
| 1 | 90 µg | 11.2 µg | 12.4 % | 200 bp |
| 2 | 72 µg | 22.3 µg | 30.9 % | 223 bp |
| 3 | 72 µg | 13.0 µg | 18.1 % | 215 bp |
| 4 | 72 µg | 8.58 µg | 11.9 % | 209 bp |

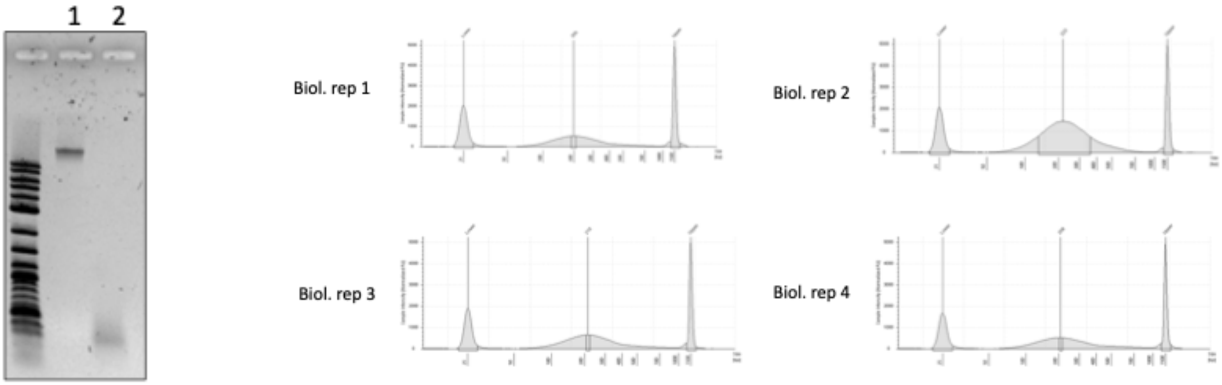


1. 10 ng/µl gDNA stock
2. Sheared DNA

**Fig. S4**. Agarose gel and tape station analysis of gDNA fragmentation.

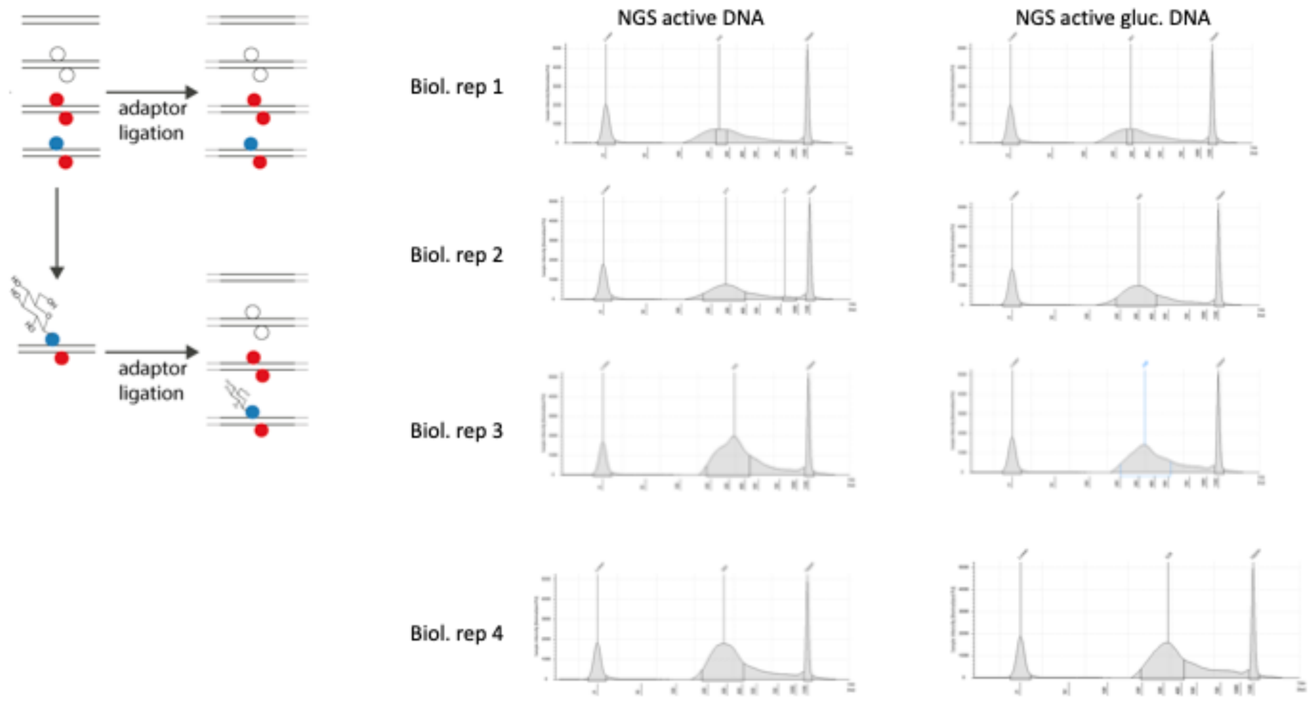


**Fig. S5**. Tape station analysis of adaptor ligation step to fragmented gDNA.

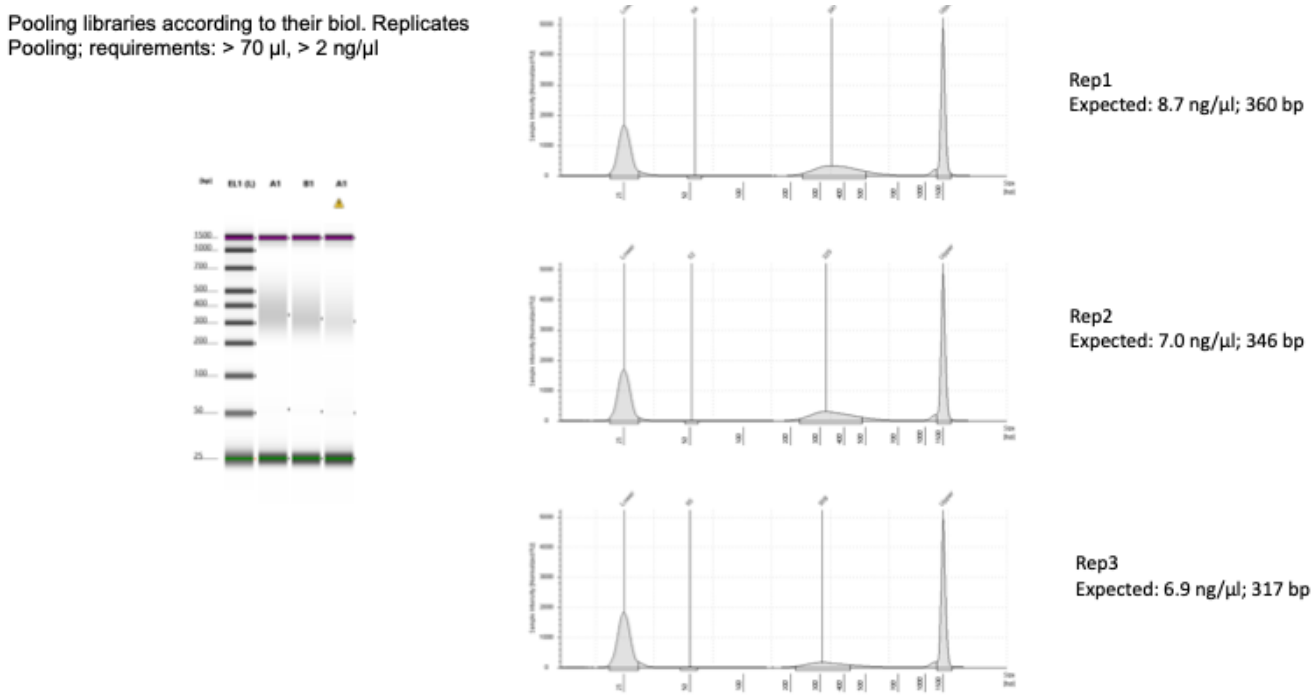


**Fig. S6**. Tape station analysis of final sequencing libraries of three biological replicates after pooling of barcoded technical replicates.

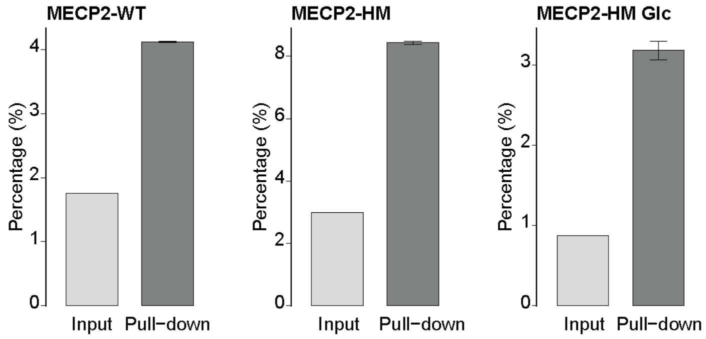


**Fig. S7**. Comparison of percentage of reads within MACS2-called peaks between input and captured samples across MECP2 conditions

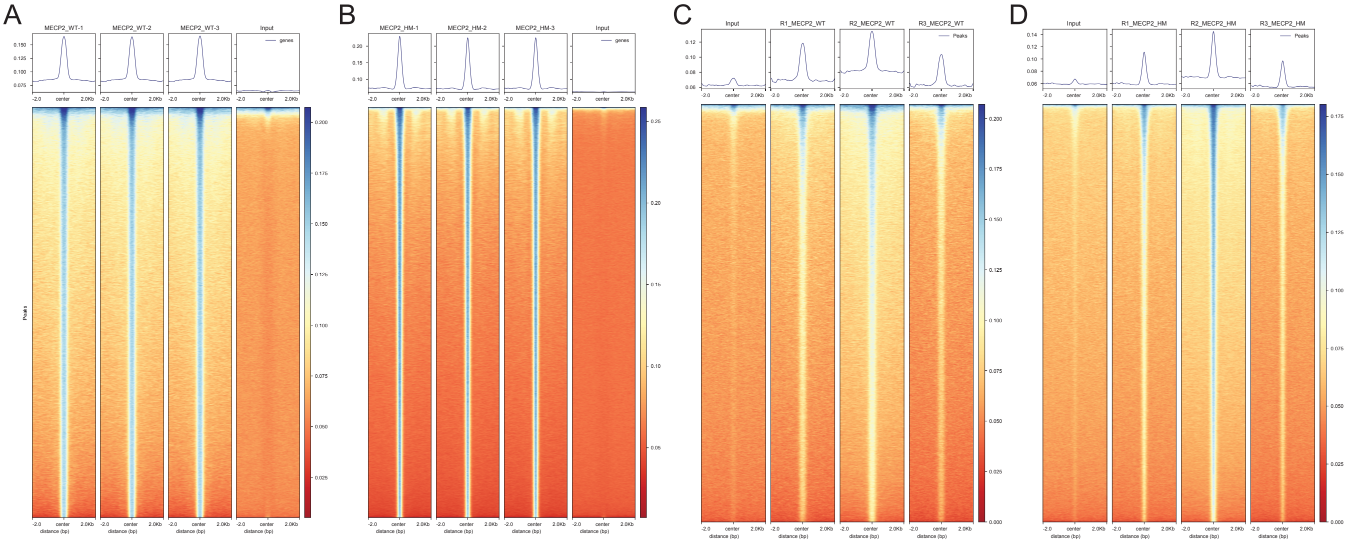


**Fig S8.** Enrichment profiles of MECP2-WT and -HM variants. Heatmaps and average signal profiles showing the enrichment of MECP2-WT (A, C) and MECP2-HM (B, D) around peak consensus regions (±2 kb from the enhancer center). (A, B) Technical replicates (Data set 1) and (C, D) biological replicates (Data set 2).

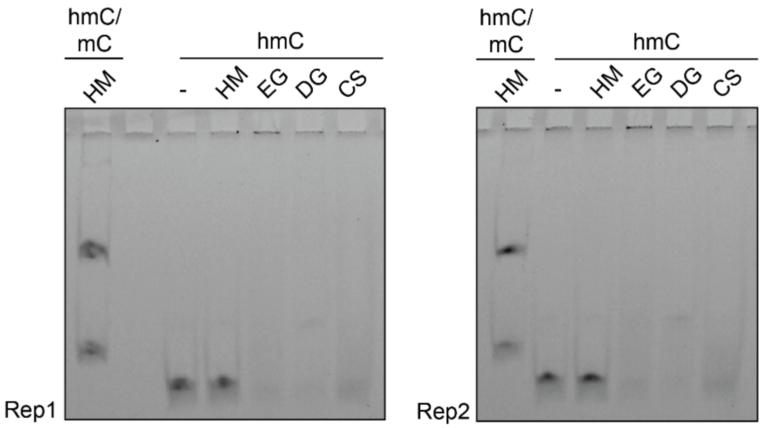


5`-6FAM-TAGGCCA**X**GTGGGAGG-3`
3`-ATCCGGTG**X**ACCCTCC-5

**Fig. S9**. Two technical replicates of electrophoretic mobility shift assays (EMSA) showing the binding of 100 nM recombinantly expressed MECP2 HM and three distinct monoclonal anti-hmC antibodies (EG – EpiGentek 0.1 mg/mL; DG – Diagenode 0.1 mg/mL; CS – Cell Signaling 0.02 mg/mL) to 750 pM of double-stranded (hmC/mC) or single-stranded (hmC) DNA.

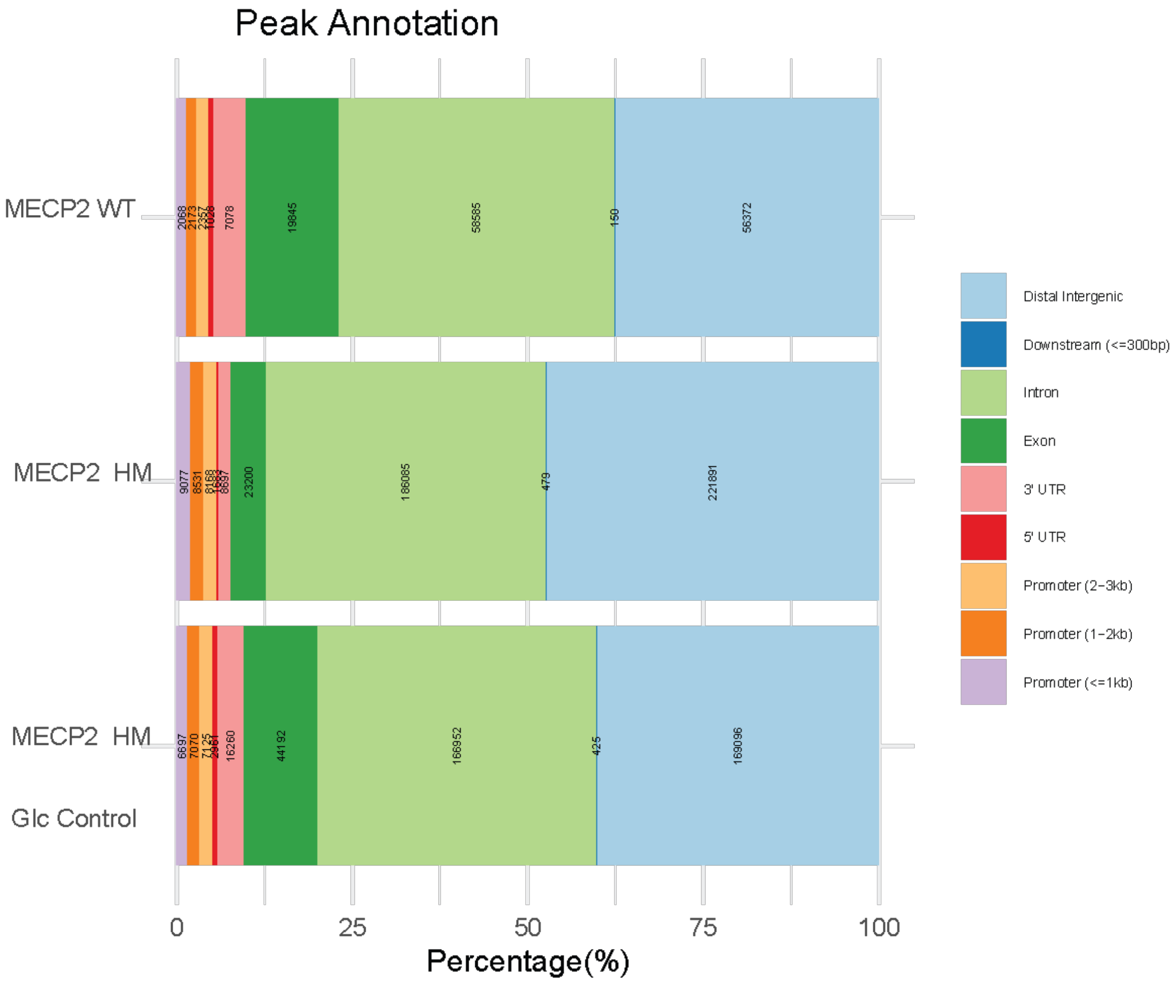


**Fig. S10**. Peak annotation profiles to genomic regions for MECP2 WT, MECP2 HM and MECP2 HM with glucosylated gDNA.

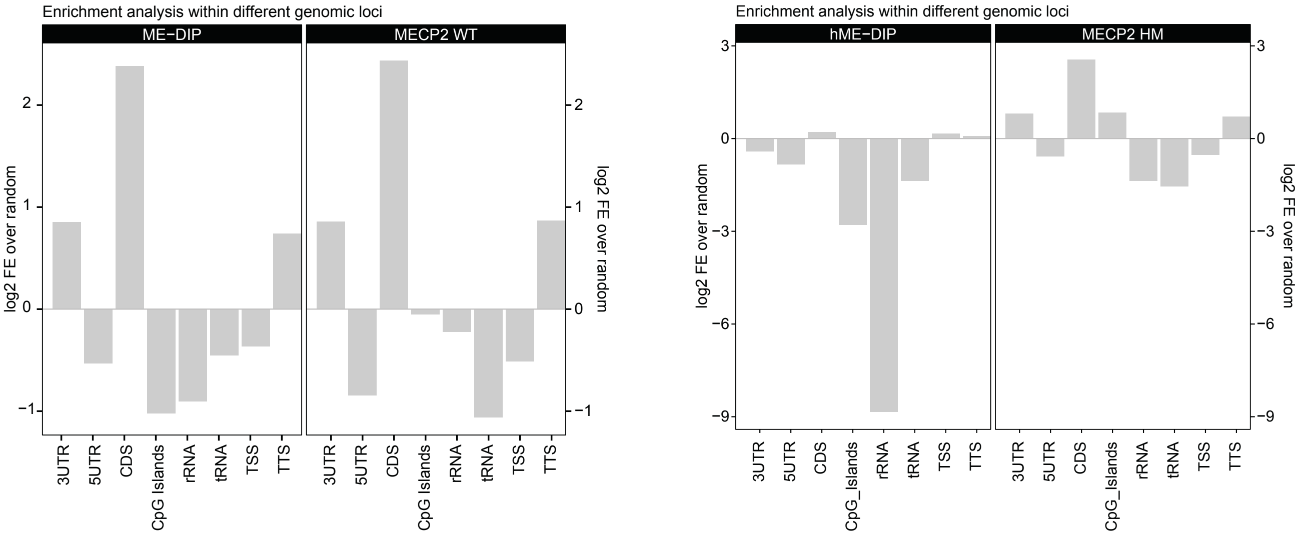


**Fig. S11**. Enrichment/depletion of selected genomic features for MeDIP and MECP2 wt as well as hMeDIP and MECP2_HM enrichments.

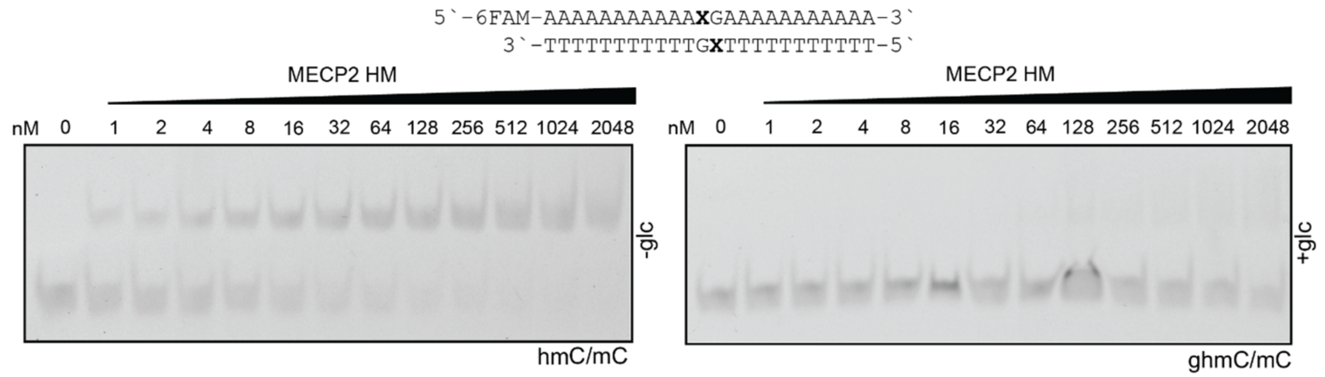


**Fig. S12**. Electrophoretic mobility shift assays (EMSA) showing the binding of MECP2 HM protein across dilution series with 2 nM of hmC/mC CpG dyad-containing 24-mer DNA duplexes before (left) and after (right) glucosylation.

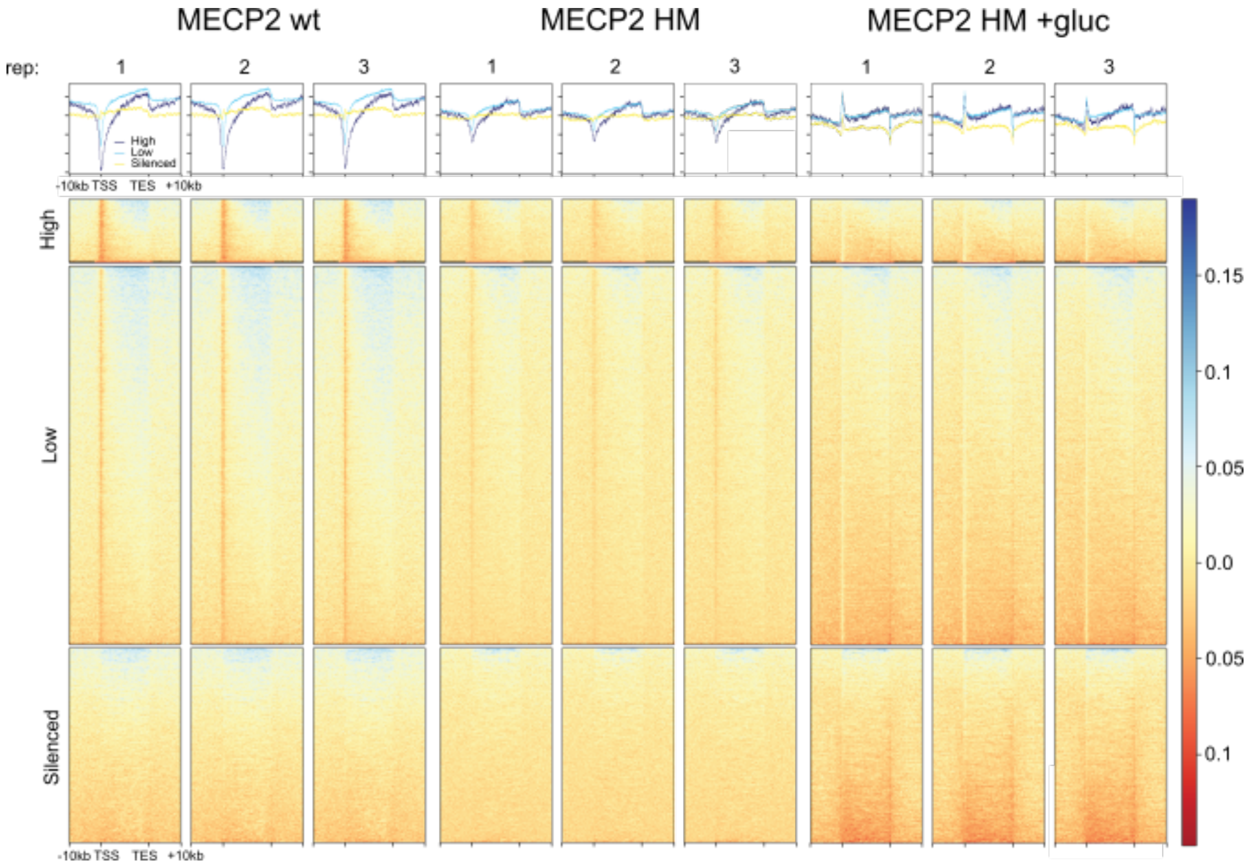


**Fig. S13**. Metagene read density profiles aligned to protein-coding genes, clustered according to their expression levels.

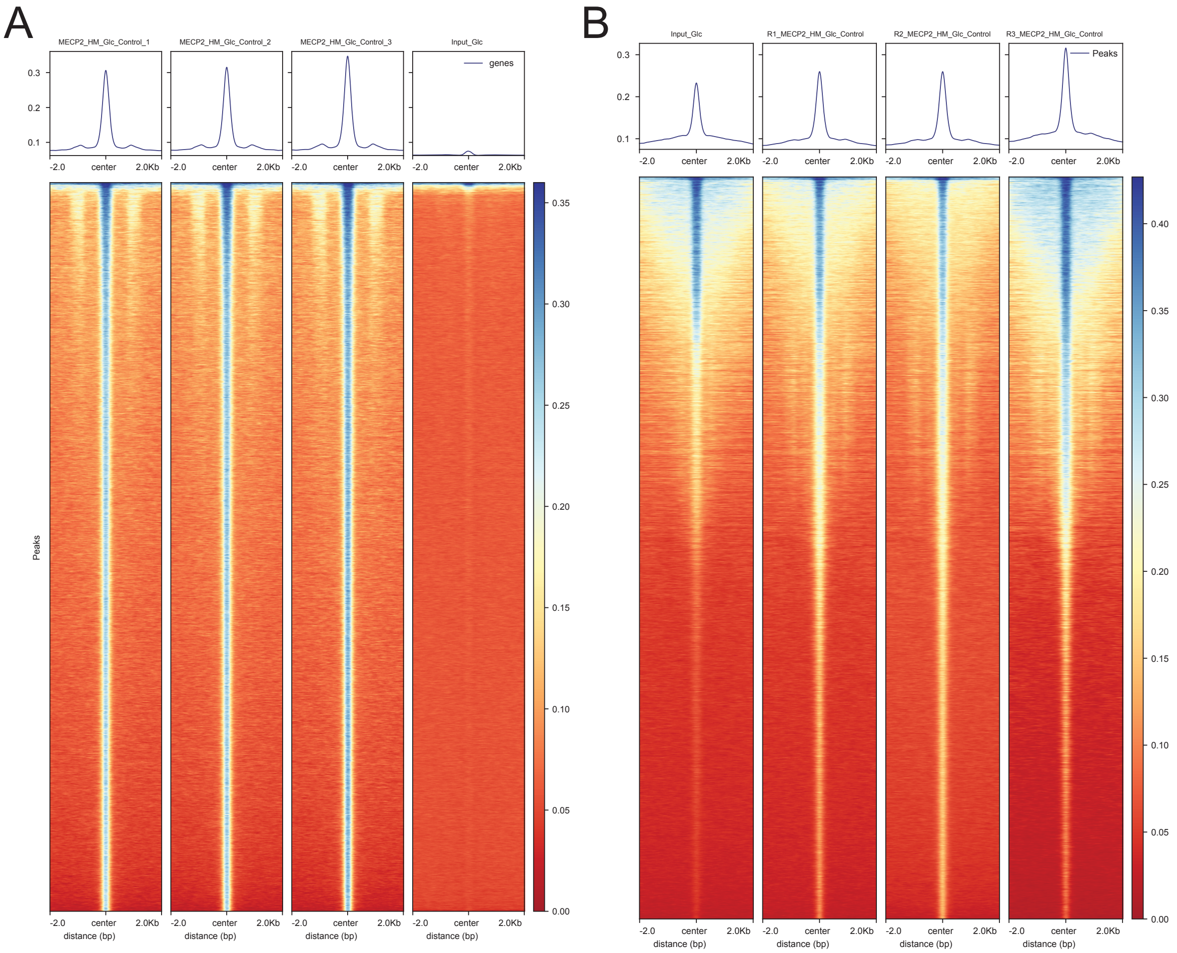


**Fig S14.** Enrichment profiles of MECP2-HM variant following glycosylation. Heatmaps and average signal profiles showing the enrichment of MECP2-HM-Glc around peak consensus regions (±2 kb from the enhancer center). (A) Technical replicates (Data set 1) and (B) biological replicates (Data set 2).

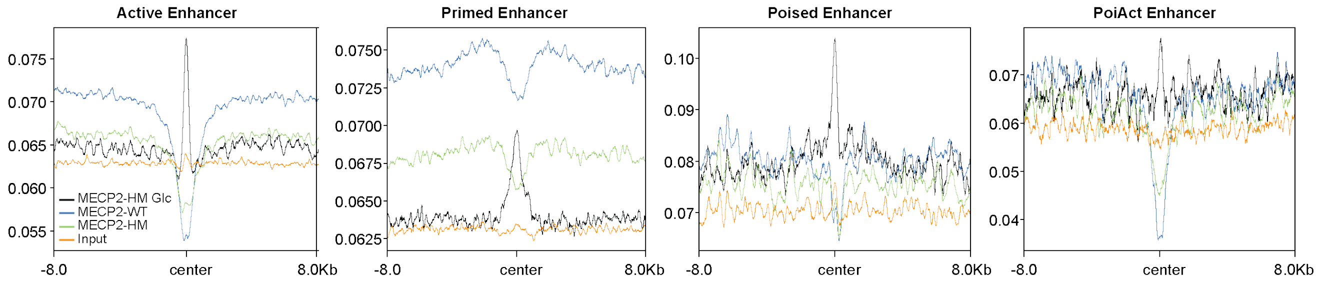


**Fig. S15**. MECP2 variant occupancies at four enhancer categories: active enhancers, primed enhancers, poised enhancers, and poised-to-active (PoiAct) enhancers.
